# Switching on the light: using metagenomic shotgun sequencing to characterize the intestinal microbiome of Atlantic cod

**DOI:** 10.1101/545889

**Authors:** Even Sannes Riiser, Thomas H.A. Haverkamp, Srinidhi Varadharajan, Ørnulf Borgan, Kjetill S. Jakobsen, Sissel Jentoft, Bastiaan Star

**Affiliations:** Centre for Ecological and Evolutionary Synthesis, Department of Biosciences, University of Oslo, PO Box 1066, Blindern, N-0316 Oslo, Norway; Department of Epidemiology, Norwegian Veterinary Institute, Oslo, Norway; Department of Mathematics, University of Oslo, PO Box 1053, Blindern, N-0316 Oslo, Norway

## Abstract

The biological roles of the intestinal microbiome and how it is impacted by environmental factors are yet to be determined in wild marine fish species. Atlantic cod (*Gadus morhua*) is an ecologically important species with a wide-spread distribution in the North Atlantic Ocean. 16S rRNA-based amplicon analyses found no geographical differentiation between the intestinal microbiome of Atlantic cod from different locations. Nevertheless, it is unclear if this lack of differentiation results from an insufficient resolution of this method to resolve fine-scaled biological complexity. Here, we take advantage of the increased resolution provided by metagenomic shotgun sequencing to investigate the intestinal microbiome of 19 adult Atlantic cod individuals from two coastal populations in Norway – located 470 km apart. Our results show that the intestinal microbiome is dominated by the *Vibrionales* order, consisting of varying abundances of *Photobacterium, Aliivibrio* and *Vibrio* species. Moreover, resolving the species community to unprecedented resolution, we identify two abundant species, *P. iliopiscarium* and *P. kishitanii,* which comprise over 50% of the classified reads. Interestingly, genomic data shows that the intestinal *P. kishitanii* strains have functionally intact *lux* genes, and its high abundance suggests that fish intestines form an important part of its ecological niche. These observations support a hypothesis that bioluminescence plays an ecological role in the marine food web. Despite our improved taxonomical resolution, we identify no geographical differences in bacterial community structure, indicating that the intestinal microbiome of these coastal cod is colonized by a limited number of closely related bacterial species with a broad geographical distribution that are well suited to thrive in this host-associated environment.

## 1. Introduction

The fish intestinal microbiome comprises a complex and specialized gut bacterial community providing a multitude of biological functions in the host, including metabolism, growth, development and immunity (reviewed in Wang et al., 2017; Ghanbari et al., 2015; Sullam et al., 2012; Izvekova et al., 2007). For instance, studies of laboratory-reared zebrafish have demonstrated that the intestinal microbiome regulates 212 genes stimulating gut epithelial proliferation, promotion of nutrient metabolism, and innate immune responses (Rawls *et al.*, 2004). Moreover, several studies of aquaculture freshwater fish have shown that gut bacterial communities produce a wide range of digestive enzymes (Sugita *et al.*, 1997; Bairagi *et al.*, 2002), and is involved in synthesis of vitamins (Sugita *et al.*, 1991). Despite this known biological importance, the composition of the intestinal microbiome in wild fish populations remains poorly understood. To date, studies of the fish intestinal microbiome have revealed a limited phylogenetic diversity, with genera from Proteobacteria, Firmicutes and Bacteroidetes constituting up to 90% of the sequence reads across different species (Verner-Jeffreys *et al.*, 2003; Ward *et al.*, 2009; Ghanbari *et al.*, 2015; Givens *et al.*, 2015; Riiser *et al.*, 2018; Talwar *et al.*, 2018). Apart from this relatively low bacterial diversity, several studies have reported a limited geographical differentiation between intestinal bacterial communities, indicating a strong influence of host-associated factors on the composition of the gut microbiome (Ye *et al.*, 2014; Llewellyn *et al.*, 2016; Riiser *et al.*, 2018). Nevertheless, most studies have been limited either because of their focus on cultured fish species (Desai *et al.*, 2012; Wu *et al.*, 2013; Zarkasi *et al.*, 2014, 2016; Schmidt *et al.*, 2016; Dehler *et al.*, 2017) or because of methodological approaches that offer limited taxonomical resolution (e.g. 16S rRNA amplicon sequencing (Star *et al.*, 2013; Ye *et al.*, 2014; Llewellyn *et al.*, 2016; Riiser *et al.*, 2018) or dependence on bacterial cultivation (Kim *et al.*, 2007; Martin-Antonio *et al.*, 2007; Valdenegro-Vega *et al.*, 2013)). Therefore, there remains a lack of detailed, baseline compositional data comparing healthy wild fish from the same species that live in different habitats with a variety of environmental conditions (Uchii *et al.*, 2006; Egerton *et al.*, 2018).

Atlantic cod (*Gadus morhua*) is an economically, ecologically and culturally important species of the North Atlantic Ocean, and represents a unique study system of the fish gut microbiome for fundamental as well as applied purposes. First, Atlantic cod, as well as the whole gadiform lineage, has lost the Major Histocompatibility Complex (MHC) II of the adaptive immune system (Star *et al.*, 2011; Malmstrøm *et al.*, 2016). This species also has an altered set of Toll-like receptors (TLRs), with a lack of TLR 1, 2, 3 and 4, and gene expansions of the intracellular TLR 7, 8 and 9 (Star *et al.*, 2011; Malmstrøm *et al.*, 2016; Solbakken *et al.*, 2016). These components of the adaptive and innate immune system are specifically involved in bacterial and viral recognition, hence likely affect the interaction between Atlantic cod and its intestinal microbiome (Star *et al.*, 2011; Star and Jentoft, 2012; Malmstrøm *et al.*, 2016; Solbakken *et al.*, 2016). Second, Atlantic cod is exposed to a variety of environmental conditions (e.g. salinity and temperature) due to its ability to exploit a wide range of ecological niches (Righton *et al.*, 2010), which in turn may influence the composition of the host microbiome. It has a large geographical distribution, which comprises various subpopulations with divergent migratory and feeding behavior (Cohen *et al.*, 1990; Godø and Michalsen, 2000; Michalsen *et al.*, 2008; Link *et al.*, 2009), and hence possibly distinctive gut microbiomes. Finally, there have been significant investments to domesticate Atlantic cod for aquaculture purposes. Various factors have prevented this industry to be profitable, for instance through difficulties in immunization of juvenile cod (Samuelsen *et al.*, 2006; Froese, Rainer and Pauly, 2012), but also through to an inefficient digestion of formulated food of larvae in the pre-stomach stage (Hamre, 2006; Lie *et al.*, 2018). Providing baseline data of the natural composition of intestinal microbiome in Atlantic cod may help efforts to improve the profitability of this industry.

The intestinal microbiome of Atlantic cod has so far been studied using both culture-based methods (Ringø *et al.*, 2006; Dhanasiri *et al.*, 2011) and culture-independent methods based on 16S rRNA amplicon sequencing (Star et al., 2013, Riiser et al. 2018). These methods show an abundance of *Bacteroidales, Erysipelotrichales, Clostridiales* and especially *Vibrionales (*Ringø *et al.*, 2006; Dhanasiri *et al.*, 2011; Star *et al.*, 2013; Riiser *et al.*, 2018). A single *Vibrionales* oligotype was found to numerically dominate the Atlantic cod intestinal microbiome, comprising more than 50% of all the sequence data (Riiser et al. 2018), suggesting that these microbiomes are not particularly complex. It is well known however, that 16S rRNA-based analyses can be confounded by amplification bias, 16S rRNA gene copy number variation and a lack of taxonomic resolution (Konstantinidis *et al.*, 2006; Liu *et al.*, 2008; Youssef *et al.*, 2009; Vasileiadis *et al.*, 2012; Shakya *et al.*, 2013; Birtel *et al.*, 2015; Amore *et al.*, 2016; Noecker *et al.*, 2016; Zhang *et al.*, 2018). It has been found that 16S rRNA has an especially low power in distinguishing various *Vibrionales* species (Sawabe *et al.*, 2007; Machado and Gram, 2015), and therefore substantial species differentiation may exist in these communities in absence of 16S rRNA divergence (Konstantinidis *et al.*, 2006; Noecker *et al.*, 2016). These limitations can be mitigated by the use of shotgun metagenomics, which offers enhanced detection of bacterial species, a better estimation of diversity, and a more in-depth insight into the functional composition of microbiomes (Llewellyn *et al.*, 2014; Romero *et al.*, 2014; Ghanbari *et al.*, 2015; Merrifield and Rodiles, 2015; Colston and Jackson, 2016; Ranjan *et al.*, 2016; Tarnecki *et al.*, 2017). Despite these advantages, however, only a handful of studies has used metagenomics approaches to investigate the intestinal microbiome in fish, and the existing studies are all limited in their number of samples investigated, their community characterization at the lower taxonomical levels (i.e. species) or geographical sampling range, with a focus on Pacific aquaculture species (Xing *et al.*, 2013; Xia *et al.*, 2014; Hennersdorf *et al.*, 2016; Tyagi *et al.*, 2019). Nevertheless, there exist no studies that use metagenomic shotgun sequencing to characterize the geographical structure and community complexity in the intestinal microbiome of wild fish.

Here, we investigate the intestinal microbial community structure of 19 adult individuals of coastal Atlantic cod from different habitats in Norway, located 470 km apart (Fig. 1a) using metagenomic shotgun sequencing. No geographical differentiation of the intestinal microbiome between these locations was previously observed based on 16S rRNA amplicon sequencing (Riiser *et al.*, 2018), providing an opportunity to test the enhanced resolution of shotgun metagenomics in a spatial and environmental context. First, we compare the genome-wide taxonomic composition and diversity based on metagenomic shotgun sequencing to that of the 16S rRNA marker-gene analysis. Second, we assess strain-level variation of the most abundant bacterial members of the intestinal community by using reference-based read mapping and comparing genome-wide single nucleotide variation. Finally, we explore the genome-wide coverage of the two most abundant bacterial strains in the Atlantic cod intestines to infer the functionality of specific genes and loss of genes.

**Figure 1:**
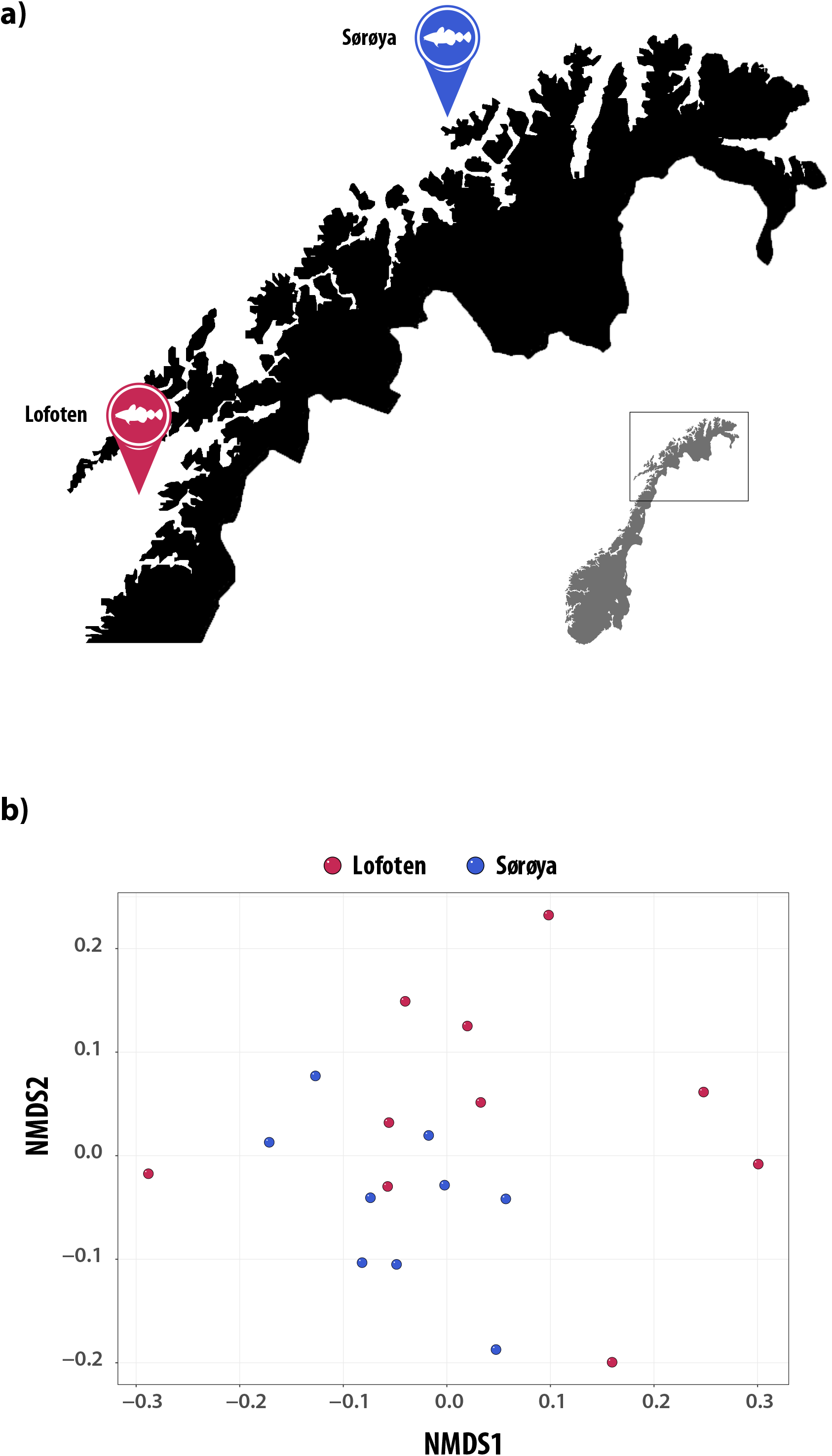
Microbial intestinal communities of wild Atlantic cod from two locations. **(A)** Map of sampling locations. **(B)** Non-metric multidimensional scaling (NMDS) plot of non-normalized, order-level sequence counts from samples from Lofoten (*red*) and Sørøya (*blue*) based on Bray-Curtis dissimilarity. The stress value of the NMDS plot is 0.22.

## 2. Methods

### 2.1 Sample collection

Wild coastal Atlantic cod (*Gadus morhua*) specimens were collected in Lofoten (N68.0619167, W13.5921667) (10 individuals, August 2014) and Sørøya (N70.760418, W21.782716) (9 individuals, September 2013) (Fig. 1a, Table S1). A 3 cm long part of the hindgut (immediately above the short, wider rectal chamber) was aseptically removed *post-mortem* by scalpel and stored on 70% ethanol. The samples were frozen (−20°C) for long-term storage. Relevant metadata such as length, weight, sex and maturity were registered. Age was determined by studying otoliths. Although different individuals were used here, these were collected on the same time and location as the fish used in a previous 16S rRNA-based study (Riiser *et al.*, 2018). We always strive to reduce the impact of our sampling needs on populations and individuals. Therefore, samples were obtained as a byproduct of conventional business practice. Specimens were caught by commercial vessels, euthanized by local fishermen and were intended for human consumption. Samples were taken post-mortem and no scientific experiments have been performed on live animals. This sampling follows the guidelines set by the “Norwegian consensus platform for replacement, reduction and refinement of animal experiments” (Norecopa) and does not fall under any specific legislation in Norway, requiring no formal ethics approval.

### 2.2 Sample preparation and DNA extraction

Intestinal samples were split open lengthwise, before the combined gut content and mucosa was gently removed using a sterile disposable spatula. Each individual sample was washed in 500 μl 100% EtOH and centrifuged before the ethanol was allowed to evaporate, after which dry weight was measured before proceeding to DNA extraction. DNA was extracted from between < 10 and 300 mg dry weight of gut content using the *MoBio Powersoil HTP 96 Soil DNA Isolation Kit (*Qiagen, Valencia, CA, USA) according to the DNA extraction protocol (v. 4.13) utilized by the Earth Microbiome Project (Gilbert *et al.*, 2010). DNA was eluted in 100 μl Elution buffer, and stored at −20° Celsius. Due to high methodological consistency between biological replicates in previous experiments, only one sample was collected per fish (Riiser *et al.*, 2018).

### 2.3 Sequence data generation and filtering

Quality and quantity of the DNA was measured using a Qubit fluorometer (Life Technologies, Carlsbad, CA, USA), and normalized by dilution. DNA libraries were prepared using the *Kapa HyperPlus* kit (Roche Sequencing, Pleasanton, CA, USA) and paired-end sequenced (2×125 base pairs) on an Illumina HiSeq2500 using the HiSeq SBS V4 chemistry with dual-indexing in two independent sequencing runs. Read qualities were assessed using *FastQC (*Andrews, 2010), before adapter removal, singleton read identification, de-duplication and further read quality trimming was performed using *Trimmomatic (*ver. 0.36) (Bolger *et al.*, 2014) and *PRINSEQ-lite (*ver. 0.20.4) (Schmieder and Edwards, 2011) (Table S2). PhiX, host and human sequences were removed by mapping reads to the phiX reference genome [GenBank:J02482.1], the Atlantic cod genome assembly (gadMor 2), (Tørresen *et al.*, 2017) and a masked version of the human genome (HG19) (Genome Reference Consortium, 2009) using *BWA (*ver. 0.7.13) (Li and Durbin, 2009) or *BBMap (*ver. 37.53) (JGI) with default parameters, and discarding matching sequences using *seqtk (*ver. 2012.11) (Li, 2012). All sequence data have been deposited in the European Nucleotide Archive (ENA) under study accession number PRJEB29346.

### 2.4 Taxonomic profiling

Taxonomic classification of quality-trimmed and filtered metagenomic paired-end reads was performed using *Kaiju (*ver. 1.5.0) (Menzel and Krogh, 2016) (“greedy” heuristic approach, -e 5), with the NCBI *nr* database (rel. 84) (incl. proteins from fungal and microbial eukaryotes) as reference (O’Leary *et al.*, 2016). Counts of reads successfully assigned to orders and species were imported into *RStudio (*ver. 1.1.383) (Racine, 2010) based on *R (*ver. 3.4.2) (R Core Team, 2017) for further processing. Final results were visualized using the R package *ggplot (*ver. 2.2.1) (Wickham, 2009). Note: Based on a recent reclassification (Machado and Gram, 2017), we refer to the reference strain *Photobacterium phosphoreum* ANT-2200 (acc. nr. GCF_000613045.2) as *Photobacterium kishitanii (*Table S3).

### 2.5 Assessment of *Vibrionales* species resolution based on 16S rRNA V4 region

RNA sequences of the most highly abundant *Vibrionales* species were downloaded from RefSeq (accessed 12.12.18) (Table S4), before 16S rRNA sequences were extracted using a custom script. Next, the 16S rRNA sequences were imported into *Geneious (*ver. 10.2.2) (Geneious), where the V4 regions (one or multiple from the same assembly) were identified and extracted. Finally, the V4 regions of the different *Vibrionales* species were aligned (File S1) using the MAFFT algorithm with default parameters, generating a sequence similarity matrix (Table S5).

### 2.6 Sequence variation analysis and genome similarity estimations

In order to assess the heterogeneity of *Vibrionales* species in our bacterial populations, we analyzed the sequence variation in *Vibrionales* genomes present in the intestinal metagenome of each fish. Initially, paired-end reads from each sample were mapped against 109 complete or scaffold-level *Photobacterium, Aliivibrio* or *Vibrio* genomes downloaded from NCBI RefSeq (rel. 84) (O’Leary *et al.*, 2016) (Table S6). The relatedness between the 15 reference genomes recruiting the highest portion of reads (Table S3) was then estimated based on Average Nucleotide Identity (ANI) and Mash genome distances using *FastANI (*ver. 1.1) (Jain *et al.*, 2018) and *Mash (*ver. 2.1) (Ondov *et al.*, 2016) (Fig. S1, Table S7). For the sequence variation analysis, paired-end reads from each individual were mapped to the 15 reference genomes using the *Snakemake* workflow (Köster and Rahmann, 2012) of *anvi’o (*ver. 5.1) (Eren *et al.*, 2015a) with default parameters in the “all-against-all” mode (with *anvi-profile--min-coverage-for-variability 10*). In *anvi’o*, contigs are divided into “splits” of maximum 20,000 bp. Splits with outlier mean coverage values (above the 98-percentile, 4-7 splits per sample), potentially containing repetitive sequences, were removed, and samples of low coverage were filtered (0-2 samples per reference genome). For each individual sample, variable sites (with min. 10X coverage) were identified, and the mean number of these per 1000 bp calculated (variation density). Next, variable sites with a minimum of 10X coverage in *all* samples were defined as single nucleotide variants (SNVs, *anvi-gen-variability-profile--min-occurrence 1--min-coverage-in-each-sample 10*). Coverage, variation density and SNV profiles were plotted in *RStudio* following the *R* script provided by *anvi’o (*Eren *et al.*, 2015b). The *anvi’o* SNV output was converted to .vcf format using a custom-developed script (https://github.com/srinidhi202/AnvioSNV_to_vcf), and the resulting .vcf files were used for principal component analysis (PCA) to test for geographical differences as implemented in *smartpca (*ver. 6.1.4) (*EIGENSOFT*) (Patterson *et al.*, 2006). The variant analysis results of six reference genomes that represent different species clusters (based on average nucleotide identity) are reported in the results section.

### 2.7 Statistical analysis

Differences in order-level classification between metagenomic shotgun sequencing and 16S rRNA amplicon sequencing (Fig. 2) was tested using ANOVA for compositional data (van den Boogaart and Tolosana-Delgado, 2013, section 5.3.3.2) using the R package *compositions (*ver. 1.40-2) (van den Boogaart and Tolosana-Delgado, 2008). Six orders common to both approaches (Fig. 2 legend, bold) and an “others” category (which contained the remaining orders) were used for the ANOVA test. Model assumptions were verified as described in section 5.3.8 of van den Boogaart and Tolosana-Delgado, 2013. Within-sample diversity (alpha diversity) was calculated using the *diversity* function in the R package *vegan (*ver. 2.4-1) (Oksanen *et al.*, 2017) based on Shannon, Simpson and Inverse Simpson indices calculated from non-normalized order-level read counts. Differences in alpha diversity were studied using linear regression. The optimal model (i.e. the model that best describes the individual diversity) was identified through a “top-down” strategy including all covariates (Table S8), except age and weight, which highly correlated with length (r = 0.78 and 0.94), and selected through *t*-tests. Model assumptions were verified through plotting of residuals. Differences in bacterial community structure (beta diversity) between Lofoten and Sørøya were visualized using non-metric multidimensional scaling (NMDS) plots based on the Bray-Curtis dissimilarity index, and tested using Permutational Multivariate Analysis of Variance (PERMANOVA) using the *metaMDS* and *adonis* functions in *vegan (*ver. 2.4-1) with both Bray-Curtis dissimilarity and Jaccard index. *Adonis* was run with 20,000 permutations. PERMANOVA assumes the multivariate dispersion in the compared groups to be homogeneous; this was verified (*p* > 0.05) using the *betadisper* function (*vegan*) (Table S9). All beta diversity analyses were based on sequence counts normalized using a common scaling procedure, following McMurdie & Holmes 2014 (McMurdie and Holmes, 2014). This method multiplies the sequence count of every unit (e.g. species) in a given library with a factor corresponding to the ratio of the smallest library size in the dataset to the library size of the sample in question, replacing rarefying (i.e. random sub-sampling to the lowest number of reads). Normalizing using this procedure effectively results in the library scaling by averaging an infinite number of repeated sub-samplings. PERMANOVA analysis was performed on normalized counts of reads classified at the order-and species level (*Kaiju*). We used Tracy-Widom and Chi-squared statistics, as implemented in *smartpca (*Patterson *et al.*, 2006), to test for significant geographical differences in the distribution of SNVs per *Vibrionales* reference genome, while correcting for multiple testing using sequential Bonferroni (Holm, 1979).

**Figure 2:**
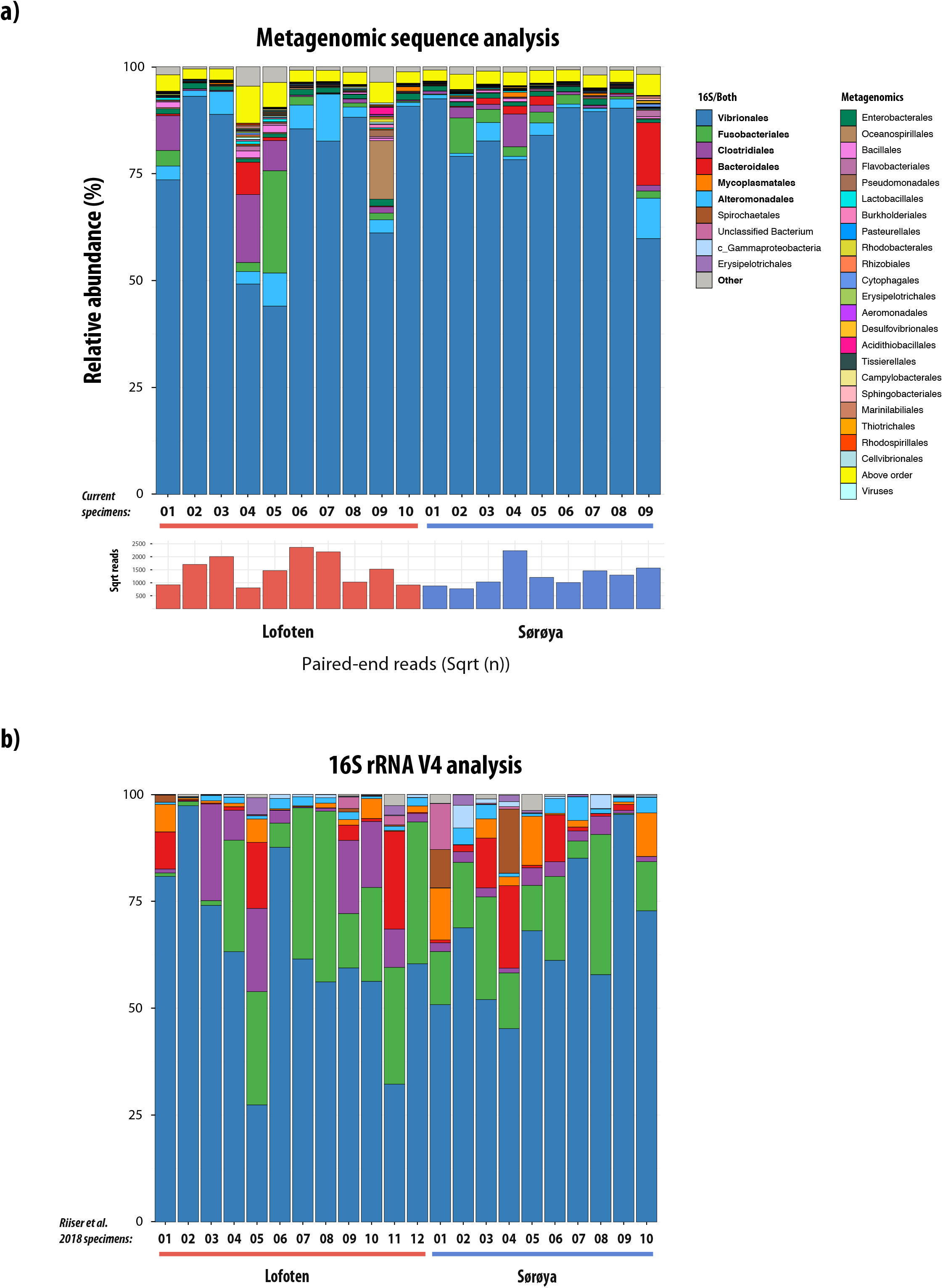
Taxonomic composition of the intestinal microbiome in Atlantic cod specimens from Lofoten and Sørøya. **(A)**Relative abundance of metagenomic shotgun sequences classified to bacterial orders. Colors represent the 30 orders with highest relative abundance, including reads that could not be assigned to the order level (*yellow*). Numbers 1-10 and 1-9 represent individual specimens. Bars below the stacked bar plot show the square root transformed counts of paired-end reads classified to order level per individual. (**B)** Relative abundance of 16S rRNA V4 sequences from Riiser et al. 2018 classified to bacterial orders. Numbers 1-12 and 1-10 represent individual specimens. Taxa in bold are identified by both methods.

### 2.8 Genome-wide characterization of *Photobacterium*

Genome-wide coverage of the two most abundant bacterial strains (*P. kishitanii* and *P. iliopiscarium*) was obtained by mapping all paired-end reads from each cod specimen toward the respective reference genomes (GCF_000613045.2, GCF_000949935.1), and visualized using the *anvi’o* command “*anvi-interactive*”. Next, “*anvi-export-gene-coverage-and-detection*” (Eren *et al.*, 2015a) was used to detect genes with zero coverage in all specimens and that are therefore consistently absent in these bacterial strains. The sequences of genes from those missing regions were used in a *blastx* search (using *blast+ (*ver. 2.6.0) (Altschul *et al.*, 1990; Camacho *et al.*, 2009)) against the *nr* database (accessed 10.12.18) using default parameters, keeping the top 5 hits. The .xml results file and gene sequences was imported into *Blast2GO (*ver. 5.2.5) (Conesa *et al.*, 2005; Conesa and Götz, 2008; Götz *et al.*, 2008, 2011), where an InterPro search (Jones *et al.*, 2014; Mitchell *et al.*, 2019), GO mapping, functional annotation and visualization was conducted with default parameters. *PHASTER (*Zhou *et al.*, 2011; Arndt *et al.*, 2016) was used to screen the *P. kishitanii* genome for the presence of prophages. The regions around the identified prophage sequences were manually inspected for the presence of other phage-associated genes (e.g. capsid heads, terminases, integrases) that could have been missed by the *PHASTER* algorithm.

The *lux* operon was not annotated in the original *P. kishitanii* RefSeq assembly (GCF_000613045.2). Therefore, the *P. kishitanii lux* genes (Table S10) were identified using the *lux* sequences of *Photobacterium phosphoreum (*AB367391.1) in a local blast search against the *P. kishitanii* reference genome with *blast+ (*ver. 2.6.0) (Altschul *et al.*, 1990; Camacho *et al.*, 2009), and manually annotated. Paired-end reads from each sample were then mapped against the annotated reference genome, and reads (.bam files) mapping to the *lux* genes were combined per location using *samtools (*ver. 1.3.1) (Li *et al.*, 2009) (“samtools merge”) to yield a consensus sequence for each location per *lux* gene. The coverage distribution and possible loss of function (due to insertions, deletions, stop codons etc.) of these *lux* gene consensus sequences was inspected using *Geneious (*ver. 10.2.2) (Geneious), *Integrative Genomics Viewer (*ver. 2.4.16) (Robinson *et al.*, 2011; Thorvaldsdóttir *et al.*, 2013) and the *ExPASy Translate* online tool (Artimo *et al.*, 2012).

## 3. Results

### 3.1 The Atlantic cod intestinal microbiome order-level composition

We obtained a dataset of 198 million paired-end reads from 19 specimens caught in the coastal waters of Lofoten (*n* =10) and Sørøya (*n* = 9, Fig. 1a). After quality trimming, the number of reads for each specimen varied from 835,000 to 7,000,000 reads (average 3 million sequences), comprised between 18.2-91.5% (mean: 66.7%) of host (Atlantic cod) DNA and between 8.5-81.8% (mean: 33.3%) bacterial DNA (Table 1, Table S11). 80% of the paired-end reads were classified, of which 96% at the order level (Table S12). The community profiles, based on non-normalized read counts, show a large overlap when clustering individuals from Lofoten and Sørøya using multivariate non-metric multidimensional scaling (NMDS, Fig. 1b). The Atlantic cod intestinal microbiome is numerically dominated by bacteria of the order *Vibrionales*, which has a mean relative abundance of 81.8% and represents > 76% of the reads in all except four individuals (Fig. 2a, Table 2). In relative abundance, this order is followed by *Alteromonadales (*3.6%), *Fusobacteriales (*3.1%), *Clostridiales (*2.9%) and *Bacteroidales (*1.7%). In total, the five orders with highest relative abundance constitute 94% of all classified sequences. A 16S rRNA-based analysis from the same locations shows that *Vibrionales* are the most abundant, followed by *Fusobacteriales, Clostridiales, Bacteroidales and Alteromonadales (*Fig. 2b, Table 2, reproduced from Riiser et al., 2018). A statistical comparison detects significant differences in the classification of the Atlantic cod intestinal microbiome comparing metagenomic shotgun sequencing to 16S rRNA-based analysis (ANOVA for compositional data, *p* = 10^-10^). In particular, the *Fusobacteriales* have a mean relative abundance of 17.1% in the 16S rRNA-based analysis vs. 3.1% in the metagenomic shotgun sequencing (Table 2). Overall, geographic location has no significant effect on the composition of the Atlantic cod intestinal microbiome (ANOVA for compositional data, *p* = 0.58) for either the metagenomic shotgun sequencing or 16S rRNA-based classification.

**Table 1:** Metagenomic sequences before and after trimming, quality filtering and host DNA removal. The table shows per sample the number of original (raw) reads, the percentage of reads remaining after trimming and filtering, percentage of host DNA, percentage of bacterial DNA and the final number of reads used in the microbiome analysis. PhiX-and human DNA sequences represent a negligible proportion, and are therefore excluded from the table. The bottom two rows show total and mean values per column. On average, 33.3% of the quality filtered reads per sample are used for microbiome analysis. For details, see Table S11.

| Sample | Raw reads | After trimming/quality filtering (%) | Host DNA (%) | Bacterial DNA (%) | Final reads |
| --- | --- | --- | --- | --- | --- |
| L_01 | 10 883 740 | 85.9 | 87.3 | 12.7 | 1 187 649 |
| L_02 | 11 140 950 | 87.9 | 62.2 | 37.8 | 3 699 538 |
| L_03 | 9 891 322 | 90.2 | 41.2 | 58.8 | 5 249 515 |
| L_04 | 10 587 865 | 86.9 | 85.2 | 14.8 | 1 364 663 |
| L_05 | 8 423 091 | 89.1 | 57.7 | 42.3 | 3 171 737 |
| L_06 | 10 879 319 | 89.6 | 30.5 | 69.5 | 6 772 948 |
| L_07 | 10 082 237 | 91.8 | 31.3 | 68.7 | 6 361 506 |
| L_08 | 9 114 703 | 87.3 | 80.5 | 19.5 | 1 549 210 |
| L_09 | 11 105 189 | 89.1 | 62.2 | 37.8 | 3 733 846 |
| L_10 | 11 140 743 | 84.7 | 86.0 | 14.0 | 1 320 875 |
| S_01 | 10 631 475 | 86.0 | 87.7 | 12.3 | 1 121 431 |
| S_02 | 11 527 589 | 85.6 | 91.5 | 8.5 | 834 564 |
| S_03 | 9 855 514 | 84.2 | 83.7 | 16.3 | 1 353 671 |
| S_04 | 9 259 707 | 92.6 | 18.2 | 81.8 | 7 018 741 |
| S_05 | 11 539 193 | 82.2 | 79.3 | 20.7 | 1 959 505 |
| S_06 | 10 272 359 | 84.8 | 85.7 | 14.3 | 1 247 744 |
| S_07 | 14 395 326 | 86.5 | 77.7 | 22.3 | 2 779 436 |
| S_08 | 9 189 209 | 87.4 | 69.7 | 30.3 | 2 431 383 |
| S_09 | 8 476 896 | 88.6 | 50.4 | 49.6 | 3 727 749 |
| Total: | 198 396 427 |  |  |  | 56 885 711 |
| Mean: | 10 441 917 | 87.4 | 66.7 | 33.3 | 2 993 985 |

**Table 2:** The 10 most abundant orders in the Atlantic cod intestinal microbiome. The table shows the ten most highly abundant orders in the metagenomic shotgun sequencing analysis, their mean, minimum and maximum relative abundance. The corresponding values from the 16S rRNA-based analysis are reproduced from Riiser et. al. (2018). NP: Not present.

| Order | Metagenomic (n = 19) |  |  | 16S rRNA (n = 22) |  |  |
| --- | --- | --- | --- | --- | --- | --- |
|  | Mean (%) | Min. (%) | Max. (%) | Mean (%) | Min. (%) | Max. (%) |
| *Vibrionales* | 81.8 | 46.7 | 95.4 | 64.3 | 27.4 | 97.3 |
| *Alteromonadales* | 3.6 | 0.7 | 11.3 | 1.8 | 0.1 | 5.5 |
| *Fusobacteriales* | 3.1 | 0.0 | 25.4 | 17.1 | 0.3 | 39.9 |
| *Clostridiales* | 2.9 | 0.1 | 17.4 | 5.5 | 0.2 | 22.6 |
| *Bacteroidales* | 1.7 | 0.0 | 15.4 | 4.6 | 0.0 | 22.9 |
| *Enterobacterales* | 1.2 | 0.8 | 1.7 | NP | - | - |
| *Oceanospirillales* | 0.9 | 0.1 | 14.4 | 0.0 | 0.0 | 0.2 |
| *Bacillales* | 0.5 | 0.1 | 1.8 | NP | - | - |
| *Mycoplasmatales* | 0.2 | 0.0 | 1.1 | 3.0 | 0.0 | 12.1 |
| *Pseudomonadales* | 0.3 | 0.1 | 1.6 | 0.0 | 0.0 | 0.5 |

The individual samples vary in diversity estimated by Shannon (H), Simpson (D) and Inverse Simpson (1/D) indices based on non-normalized order-level read counts (Fig. S2, Table S13). The variation in alpha diversity is reflected in the abundance profile in Fig. 2a, where in particular, four Lofoten samples (01, 04, 05, 09) and one Sørøya sample (09) contain higher relative abundances of orders other than *Vibrionales*. Top-down reduction of linear regression models based on the alpha diversity indices ends up with models containing no significant covariates (Table S8), indicating that neither location, length or sex have an impact on the within-sample diversity. Similarly, PERMANOVA analysis based on the beta diversity measures Bray-Curtis and Jaccard reveals no statistically significant differences in community structure at the order level between Lofoten and Sørøya (Table 3).

**Table 3:**
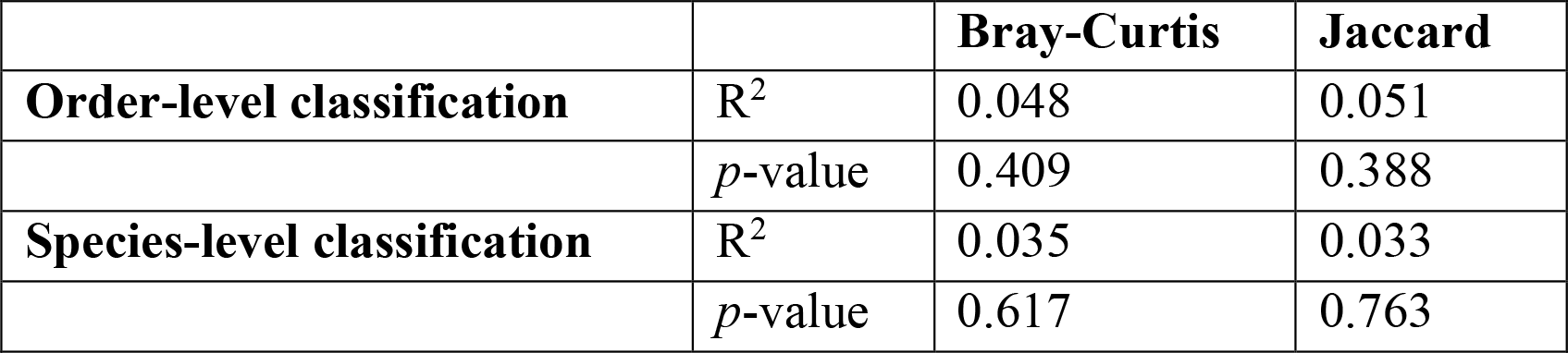
PERMANOVA analysis of diversity differences between bacterial communities from Lofoten and Sørøya (beta diversity). The table shows R^2^ and *p*-values from multivariate statistical analyses to test for community composition differences based on reads classified at order-and species level. The results are based on read counts normalized by common scaling. Degrees of freedom (*df*): 18.

### 3.2 The species-level composition within *Vibrionales*

Overall, 55.3% of the reads are classified to the species level (Table S12). Of these, *Photobacterium, Aliivibrio* and *Vibrio* species are consistently found in all individuals, and constitute between 39-94% (mean: 77.3%) of all species-level reads (Fig. 3a). The *Vibrionales* community is dominated by *P. iliopiscarium (*mean relative abundance: 40.3%) and *P. kishitanii (*MRA: 26.6%) (Fig. 3b), while specific samples also have a high relative abundance of *A. logei (*maximum relative abundance (MRA): 19.4%), *P. piscicola (*MRA: 38.8%), *A. wodanis (*MRA: 18.5%), *A. fischeri (*MRA: 10.1%) and *A. salmonicida (*MRA: 10.0%). We detect no significant difference in the intestinal *Vibrionales* species community structure between Lofoten and Sørøya (Table 3).

**Figure 3:**
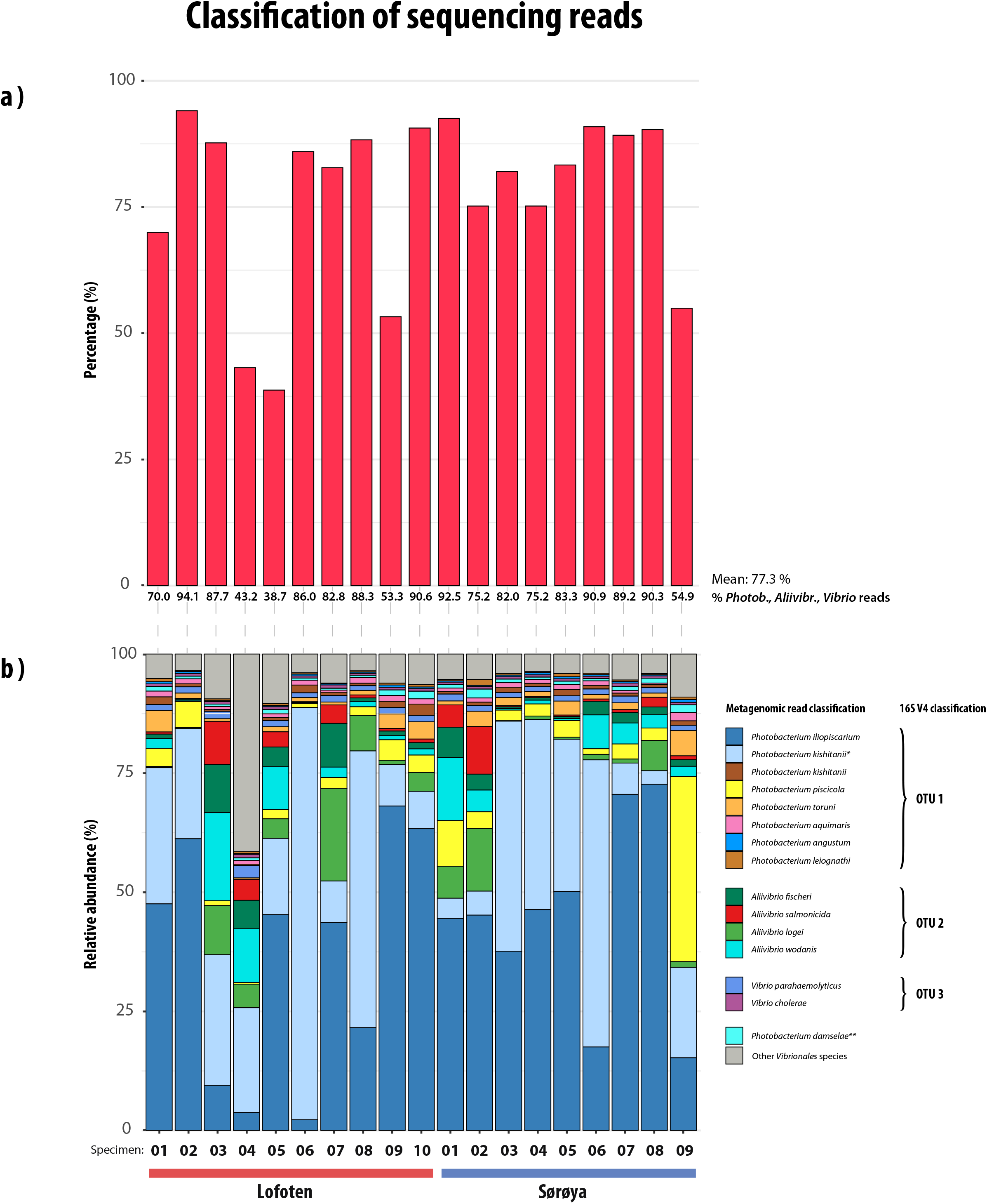
Diversity of species within *Vibrionales*. **(A)** Proportion of reads per sample classified as either *Photobacterium, Aliivibrio* or *Vibrio* species to all reads classified to species level. Percentages are shown along the x-axis. (**B)** Relative abundance of *Photobacterium, Aliivibrio* and *Vibrio* species in the Atlantic cod intestinal microbiome, as determined by protein-level classification of paired-end sequences. Colors represent the 15 species with highest relative abundance, and numbers 1-10 and 1-9 represent individual specimens. The legend is ordered by OTU membership based on clustering of the species’ 16S rRNA V4 sequences at a 97% sequence similarity level. \**Photobacterium kishitanii* strain previously classified as *Photobacterium phosphoreum* strain ANT-2200. **No V4 sequence of sufficient length available.

Metagenomic shotgun sequencing identifies a set of clearly separated, highly abundant *Photobacterium, Aliivibrio* and *Vibrio* species in the Atlantic cod intestines (Fig. 3b). We retrospectively assessed whether 16S rRNA-based taxonomic profiling is able to provide an equally detailed description of the bacterial community by analyzing the 16S V4 sequences of these *Vibrionales* species (Table S4, Fig. S3, File S1). Several of the species share identical V4 sequences (Table S5), and based on 97% sequence identity –the most frequently used parameter in 16S rRNA-based taxonomical analysis– the 14 species group into three operational taxonomic units (OTUs) (Fig. 3b). In particular, the two most highly abundant *Vibrionales* species, *Photobacterium iliopiscarium* and *Photobacterium kishitanii,* share identical V4 sequences together with five other *Photobacterium* species (Table S5).

### 3.3 Within-*Vibrionales* levels of Single Nucleotide Variant heterogeneity

We assessed the heterogeneity of the reads mapping to each of the 15 most abundant *Vibrionales* bacterial reference genomes (Table S3). These 15 genomes all obtained sufficient coverage across the majority of samples to confidently identify SNVs with a greater than 10-fold coverage. Sequence similarity estimations based on the average nucleotide identity (ANI) and mash distance among these 15 genomes reveal a clear separation between the *Aliivibrio-, Photobacterium-* and *Vibrio* species (Fig. S1, Table S7). The *Aliivibrio* species are more similar to each other than the *Photobacterium* species, and *Vibrio renipiscarium* has a higher sequence divergence compared to the other genomes. The overall differences in sequence diversity among the species (Fig. S1) are reflected in the results of SNV analysis (Fig. S4), e.g., species from the *Aliivibrio* cluster all have a lower SNV density than most *Photobacterium* species. Based on sequence similarity (%ANI) of these genomes, the results of six reference genomes that represent different species clusters are reported here (Fig. 4 and Fig. S1).

**Figure 4:**
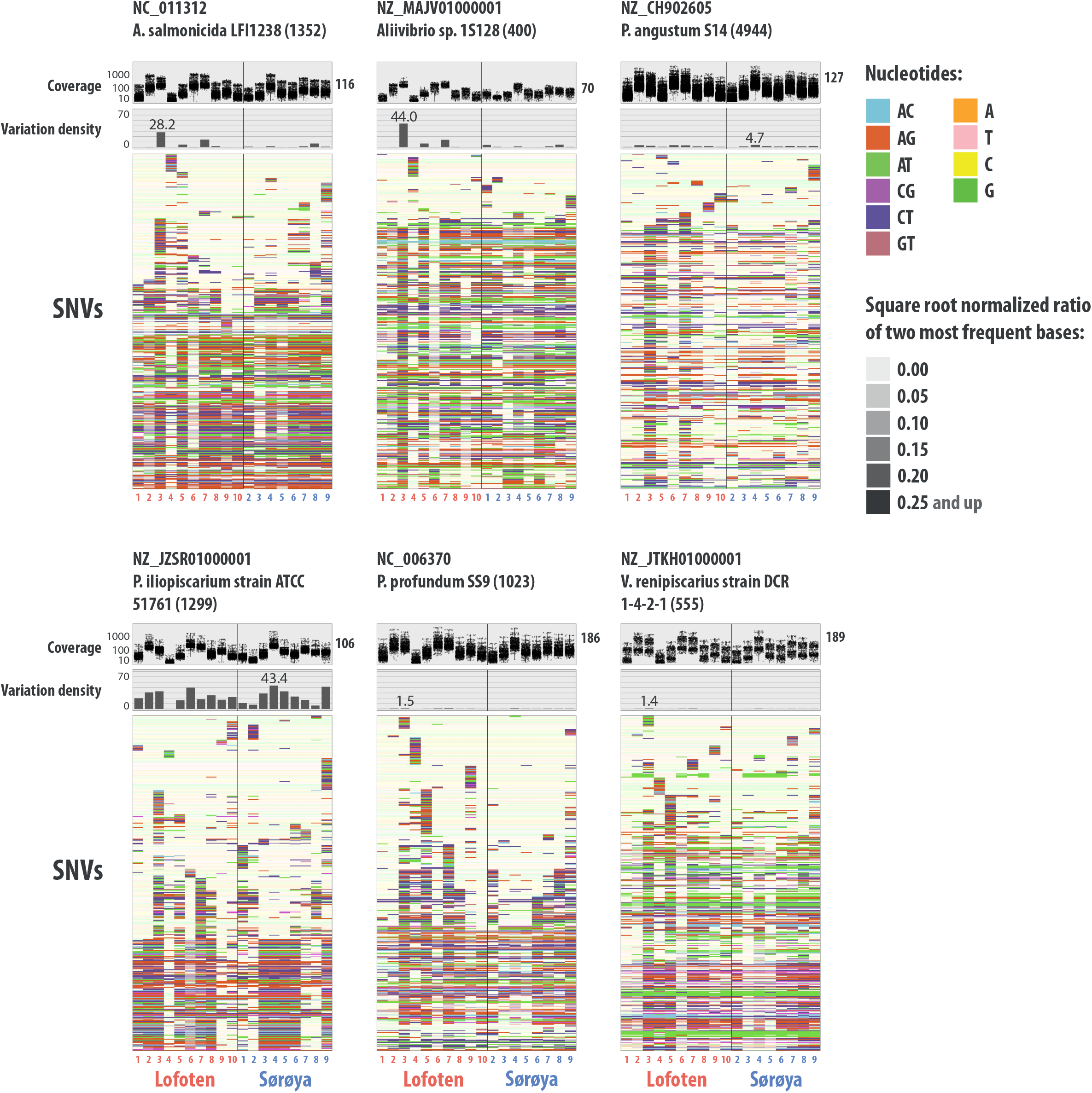
Variation analysis of *Vibrionales* reference genomes. For six *Vibrionales* reference genomes, the figure displays (from top to bottom) (1) read coverage per single nucleotide variant (SNV) position in each sample from Lofoten (*red numbers*) and Sørøya (*blue numbers*), (2) variation density (number of variable positions per kbp. reported in each individual sample, independent of coverage in the other samples) per sample and (3) heatmap of a randomly chosen subset of 400 SNVs. In the heatmap, each row represents a unique variable nucleotide position, where the color of each tile represents the two most frequent competing nucleotides in that position. The shade of each tile represents the square root-normalized ratio of the most frequent two bases at that position (i.e., the more variation in a nucleotide position, the less pale the tile is). The y-axis of the coverage-and variation density plots are scaled across the reference genomes. For each genome, the density plot (on top) is annotated with the maximum variation density value (*grey number*).

Overall, the different reference assemblies vary in the mean fold coverage, the density of variable sites within each individual sample and in the total number of SNVs observed in all samples. For instance, almost 5000 SNVs are detected in the *P. angustum* S14 genome, but the average density within specimens is low (max. 4.7/Kbp). In contrast, *P. iliopiscarium* yields less SNVs (1299) overall, yet a higher average density (max. 43.4/Kbp). The density of variable sites varies across specimens for several of the reference genomes, reflecting varying levels of heterogeneity in the bacterial populations within specimens. This pattern is particularly strong for *P. iliopiscarium,* varying from 0.1 to 43.4 variant positions per Kbp per individual specimen (Fig. 4). Likewise, the variation analysis of the two *Aliivibrio* genomes (*A. salmonicida* and *A.* sp. 1S128) indicate that sample L_03 consists of a complex mix of *Aliivibrio* strains. Despite the overall differences in SNV abundance between reference strains, we observe no statistically significant differences (based on Tracy-Widom and Chi-squared statistics) in SNV profiles between Lofoten and Sørøya among any of the 15 *Vibrionales* strains (Fig. S5, Table S14).

### 3.4 Genome-wide discrepancies between abundant *Photobacterium* strains and their closest relatives

Per individual, 85% of the *Photobacterium iliopiscarium* genome and 45% of the *Photobacterium kishitanii* genome are sequenced to a depth of minimum 5-fold coverage, respectively (Table S15). Whereas reads aligned to *P. iliopiscarium* provide near complete coverage of the entire assembly in all individuals, reads aligned to *P. kishitanii* show consistent lack of alignments in a specific genomic region between 60 and 80 kbp (Fig. 5). This region in the Mediterranean *P. kishitanii* reference genome (Table S3) contains a prophage (Machado and Gram, 2017), and the deletion found here suggests that the North Atlantic population of this species lacks this particular prophage (30 – 50 kbp), as well as other host DNA. The difference in observed coverage between the two species translates directly to the number of genes lost; while only seven genes are absent in the Atlantic cod *P. iliopiscarium* strains, 698 genes are absent (with zero coverage) in the Atlantic cod *P. kishitanii* strains compared to their reference assemblies (Fig. 5, Table S16).

**Figure 5:**
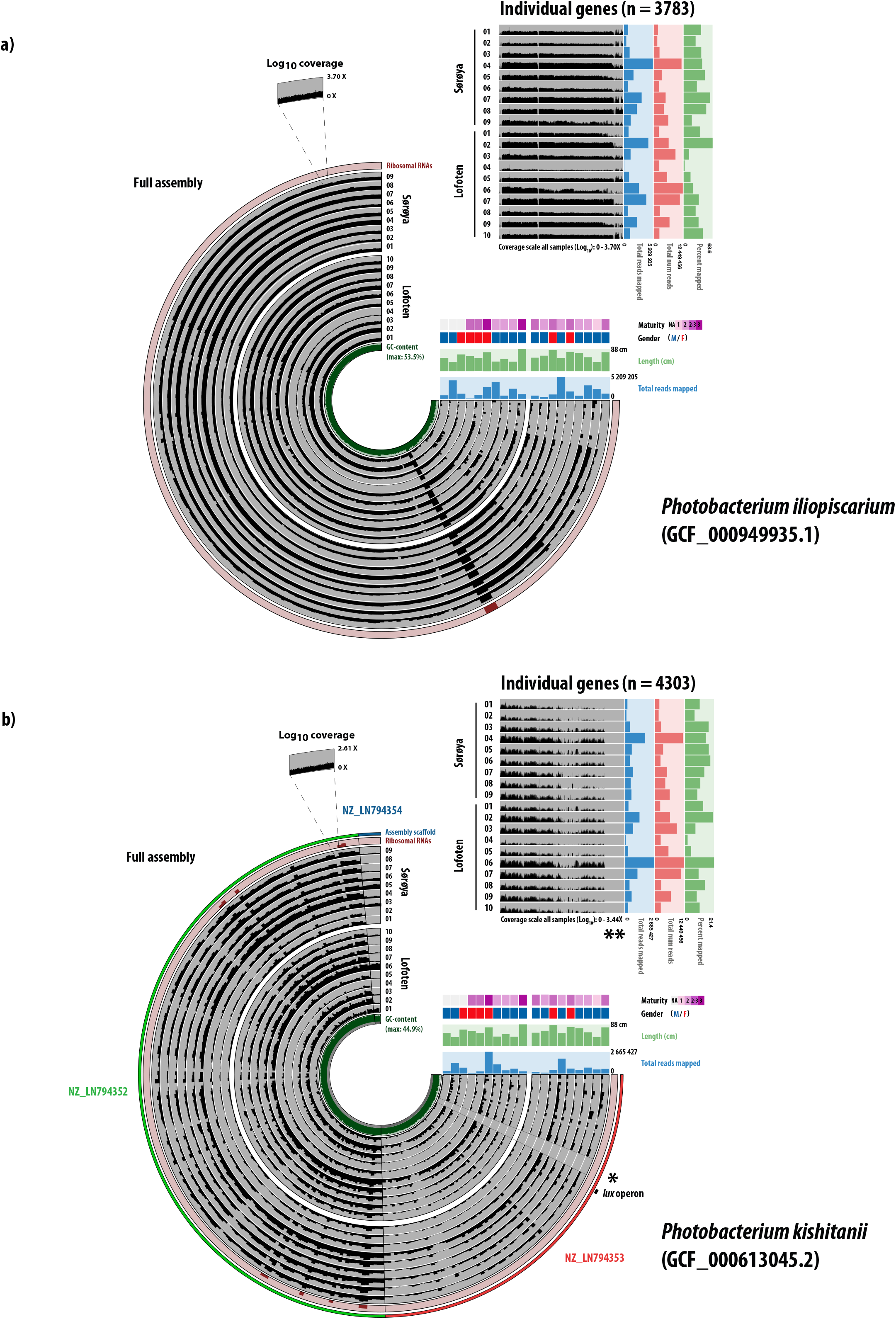
Representation of the two most abundant *Photobacterium* species among the individual samples. For each of the two most abundant *Photobacterium* species in the Atlantic cod samples, *Photobacterium iliopiscarium (***A)** and *Photobacterium kishitanii (***B)**, the figure gives an overview of the sequence coverage distribution at the assembly level (circle) and gene level (upper right square). In the assembly overview, each bar represents 20,000 bp. of a contig. Starting from the center, the concentric rings display the GC %, log10 coverage of the 19 samples, and predicted ribosomal RNAs. The coverage scale is identical for all samples, and the maximum value is given in the extracted selection above the assembly overview. The assembly overview metadata shows information on the total number of reads mapped per sample, and physical parameters associated with each individual fish. In the gene overview, each bar represents an individual gene, and the genes are ordered by differential coverage across samples. Maximum log10 coverage is given below the figure. The metadata gives information of the total number of reads and the number and percentage of reads mapped. In contrast to the incomplete *P. iliopiscarium* assembly (289 contigs) (A), the *P. kishitanii* assembly (B) consists of only three scaffolds, annotated in the outer layer. For *P. kishitanii*, the presence of the *lux* operon is annotated with a black square. The (*) denotes sections of the reference genome completely missing in *P. kishitanii* in the Atlantic cod intestines, while (**) denotes genes in the reference genome absent in the *P. kishitanii* in the cod samples.

We obtained gene ontology data for 400 of the 698 genes that are absent from the *P. kishitanii* strain (Table S17, Fig. S6). A striking number of sequences encodes membrane or membrane-associated cellular components (GO CC classification: membrane, membrane part, Fig. S6). Independent of functional annotation, a *blast* search indicate that the majority of the 698 gene sequence reads matches *P. kishitanii (*Fig. S6), confirming the presence of this species (and not *P. phosphoreum* ANT-2200, as the reference is classified) in the Atlantic cod intestines. In contrast to *P. kishitanii*, only seven genes are absent in the Atlantic cod-associated *P. iliopiscarium* strain compared to its closest relative. Only one of these is successfully annotated, and is assigned a function in “chromosome partitioning”.

### 3.5 The *Photobacterium kishitanii* lux operon

*Photobacterium kishitanii* is known to contain the *lux* operon (i.e. encoding luciferase activity) necessary for bioluminescence. Due to the high relative abundance of this bacterium (or a closely related strain) in the Atlantic cod gut, and the unclear role of such a bioluminescence feature in the intestinal compartment, we investigated the putative loss of the *lux* operon in the *P. kishitanii* strain associated with the Atlantic cod intestinal samples. All *lux* genes (*luxC, D, A, B, F, E, G*) of the operon are identified in the *P. kishitanii* strain in Atlantic cod (Fig. S7, Table S10). Their mean coverage across all samples ranges from 12.5-21X, and the coverage of each gene per sample correlates with the total number of mapped paired reads per sample (Table S10). No insertions or deletions are observed in the *lux* operon gene sequences (File S2). We find between 5-23 nonsynonymous substitutions in Atlantic cod *P. kishitanii lux* genes compared to the reference sequence. None of these substitutions results in a stop-codon, and there is no indication that the translation of the complete *lux* operon is disabled in the *P. kishitanii* strains in the Atlantic cod intestine.

## 4. Discussion

Here, we have used metagenomic shotgun sequencing to provide a first in-depth characterization of the Atlantic cod intestinal microbiome determined down to species-and strain level resolutions. In contrast to previous 16S rRNA data, which yielded a single numerically dominant OTU belonging to genus *Photobacterium (*Riiser *et al.*, 2018), we find at least nine bacterial *Photobacterium* species that occur in varying abundances in the Atlantic cod gut. Based on their 16S V4 sequences, eight of these species cluster into a single OTU (at 97% sequence identity), demonstrating the increased taxonomical resolution provided by metagenomic shotgun sequencing.

Two related species (*P. iliopiscarium* and *P. kishitanii*) are particularly abundant, comprising 67% of reads classified to genus *Photobacterium* and more than 50% of all reads classified. Both have previously been isolated from the intestines of Atlantic cod (e.g., Dhanasiri et al., 2011), although these species differ in their perceived ecological niches. *P. kishitanii* is a cosmopolitan, wide-spread facultative psychrophilic bacterium (Urbanczyk *et al.*, 2011; Machado and Gram, 2017). It is most known for containing the *lux-rib* operon, which is essential for quorum sensing and generating bioluminescence in the light organs in –amongst others– *Gadiform* deep-water fish (Ast and Dunlap, 2005). In contrast, *P. iliopiscarium* is a non-luminous bacterium that has been isolated from the intestines of several cold-water species, including Atlantic cod, yet the ecological distribution of this bacterium is still poorly known (Onarheim and Raa, 1990; Onarheim *et al.*, 1994; Urakawa *et al.*, 1999; Ast and Dunlap, 2005; Smith *et al.*, 2007). Based on phylogenetic analyses, *P. iliopiscarium* has lost the *lux-rib* operon, presumably due to niche specialization (Machado and Gram, 2017). The high abundance of *P. kishitanii* –with full repertoire of *lux* genes– in the Atlantic cod intestinal samples is particularly interesting. Zooplankton feeding on luminescent bacteria have been found to glow, which makes them more vulnerable to predation (Zarubin *et al.*, 2012). Bioluminescence has therefore been suggested to be an adaptation encouraging fish ingestion, allowing efficient dispersal of the bacteria through their fish hosts (Takemura *et al.*, 2014). Although luminescent bacteria have been long known from excrement pellets (Andrews *et al.*, 1984) and a wide range of fish taxa (Ruby and Morin, 1979), for instance captive Atlantic halibut (*Hippoglossus hippoglossus*) (Verner-Jeffreys *et al.*, 2003), their relative abundances in wild fish intestines have never been reported. Here, we observe that such luminescent bacteria comprise an abundant component (26.6% of reads for *P. kishitanii*) of the intestinal microbiota in Atlantic cod. This observation suggests that fish intestines form a particularly rich niche for bioluminescent bacteria.

We compared the genomic organization of the two most abundant *Photobacterium* species in the Atlantic cod intestinal microbiome to their closest relatives by investigating genome-wide alignments. A near complete read coverage across the reference genome of *P. iliopiscarium* was observed, indicative of limited large-scale genomic rearrangements in those strains sampled from the Atlantic cod intestines. In contrast, a consistent lack of read coverage in distinct genomic regions across the *P. kishitanii* reference genome demonstrates absence of specific regions in all Atlantic cod-associated strains. This lack results in the absence of nearly 700 of the 4300 genes annotated on the *P. kishitanii* reference assembly. Such an observation is not uncommon among *Photobacterium* species, and as little as 25% of genes is expected to be conserved between different strains of this genus (Machado and Gram, 2017). Nevertheless, the consistent absence of the same genomic region in all individuals indicates that these intestines have been colonized by a closely related *Photobacterium kishitanii* strain in both geographical locations. Interestingly, the missing genes predominantly encode components of the cell membrane. Given that the bacterial cell membrane plays a central role in host-microbiome interaction, and the fact that Atlantic cod has lost the MHC II pathway and possess a special TLR repertoire (Star *et al.*, 2011; Solbakken *et al.*, 2016), it is possible that the loss of these genes represents a functional adaptation to the peculiar immune host environment.

The functional role of *P. iliopiscarium, P. kishitanii* and other members of the genus *Photobacterium* in the Atlantic cod intestines, and the reason for their high abundance (host-selection or environmental exposure), remains unclear. Members of the *Photobacterium* genus have been shown to aid in the digestive process of Dover sole (*Solea solea*), i.e. by degrading chitin (MacDonald *et al.*, 1986), while others show antagonistic activity towards common bacterial pathogens in Atlantic cod (MacDonald *et al.*, 1986; Caipang *et al.*, 2010; Ray *et al.*, 2012; Egerton *et al.*, 2018). Such roles in protective immunity or digestion suggest an evolutionary benefit of host selection for the colonization by *Photobacterium*. Host selection for certain taxa (classified based on 16S rRNA) has been observed in zebrafish and Atlantic salmon parr (Roeselers *et al.*, 2011; Dehler *et al.*, 2017). It may be assumed that bacteria more intimately associated with their host (i.e. through a strong association with the mucosal layer relative to the general gut content) are actively selected for. Based on such an assumption, host selection for *Photobacterium* in Atlantic cod is implied by a significantly higher abundance of this genus associated with the intestinal mucosal layer relative to the gut content based on 16S RNA classification (Riiser *et al.*, 2018). Nonetheless, it is currently not clear if this higher abundance of the genus in the mucosal layer is due to the increased selection for specific *Photobacterium* strains, e.g. *P. iliopiscarium* or *P. kishitanii*. Hence, more elaborate functional studies are required to investigate the roles of *P. iliopiscarium, P. kishitanii* and the other members of *Photobacterium* in the Atlantic cod intestines, and whether their high abundance are due to its unique immune system or by external, ecological factors (Star *et al.*, 2011; Star and Jentoft, 2012).

Our results shed light on the order-level classification based on 16S rRNA amplicon sequencing versus metagenomic shotgun sequencing. There are significant differences in the order-level bacterial community composition detected by the two analysis methods. For instance, *Fusobacteriales* has an average relative abundance of 17.1% based on 16S rRNA, yet comprises 3.1% of the metagenomic shotgun data. Interestingly, a member of the *Fusobacteriales (Cetobacterium somerae*) that has been isolated from the intestinal tract of fish (Tsuchiya *et al.*, 2008), has a particularly low GC content (28.5%) (ecogenomic.org, 2013). A bias against such low GC content has been observed during library preparation (for instance due to the enzymatic fragmentation applied in our protocol), amplification and sequencing (Benjamini and Speed, 2012), and could explain lower *Fusobacteriales* relative abundance in the metagenomic data. This lower proportion of *Fusobacteriales* may contribute to the increased relative abundance of *Vibrionales* in the metagenomic shotgun data. Despite such differences however, both methods do identify a similar set of abundant microbial taxa, and show a dominant presence of *Vibrionales* in the intestines of Atlantic cod.

Several 16S rRNA-based studies have reported limited effects of geographic location on the composition and diversity of the fish intestinal microbiome. In Atlantic salmon (*Salmo salaris*), little differentiation was observed in populations from both sides of the Atlantic Ocean, and the intestinal microbial community composition was rather associated with life stage (Llewellyn *et al.*, 2016). Similarly, the gut microbiome of invasive Silver carp (*Hypophthalmichthys molitrix*) collected at highly separated sampling spots in the Mississippi river basin was affected by sampling time rather than location (Ye *et al.*, 2014). Finally, no significant differences in intestinal microbiome composition was detected in Atlantic cod from Lofoten and Sørøya, separated by 470 km (same locations as in this study) using 16S rRNA analyses (Riiser *et al.*, 2018). Our in-depth characterization of the Atlantic cod intestinal microbiome using metagenomic shotgun sequencing allowed us to re-address if significant geographical population structure could be demonstrated at the species or within-species level based on genome-wide data. First, at the level of species, we observe no significant geographical differences using genome-wide protein-based analyses. This lack of differentiation is partly due to the presence of the two *Photobacterium* species (*P. iliopiscarium* and *P. kishitanii),* which are abundant in all specimens. Second, based on SNV variation across the genome of the 15 most abundant *Vibrionales* species, we find that the gut of each fish specimen contains a unique and diverse set of strains of each species, nonetheless, no significant geographical differences are observed. Both the protein-based and strain-level approaches assessing the diversity of *Vibrionales* indicate that the microbial community composition of the gut is not related to the geographic location where the cod specimens were caught. This absence of geographical substructure, even based on genome-wide data, suggests that the intestinal microbiome of Atlantic cod is colonized by a diversity of *Vibrionales* species with a large spatial distribution.

## 5. Conclusions

We here present the first characterization of the intestinal microbiome of wild Atlantic cod using genome-wide shotgun data. Based on improved resolution, we find that two closely related *Photobacterium* species (*P. iliopiscarium* and *P. kishitanii*) are particularly abundant in the intestinal communities of Atlantic cod, comprising the majority of reads. Interestingly, our results show that luminescent bacteria comprise an abundant component of the intestinal microbiota in Atlantic cod. Notwithstanding our improved taxonomical resolution, no significant differentiation at the species or within-species level between Lofoten and Sørøya was detected, indicating that the composition of the intestinal microbiome is not related to the geographic location of the Atlantic cod specimens.

## Supporting information

16S V4 sequences

lux gene sequences

## Authors’ contributions

SJ, BS and THA conceived and designed the experiments. KSJ provided the initial framework for the study. ESR and SJ sampled the specimens. ESR performed the laboratory work. ESR and THA performed data analysis. SV created the Python script to convert the *anvi’o* format to .vcf. ØB, THA, ESR and BS interpreted the results. ESR and BS wrote the paper with input of all authors. All authors read and approved the final manuscript.

## Acknowledgements

We thank Børge Iversen and Helle Tessand Baalsrud for their kind help in sampling cod specimens in Lofoten, and Martin Malmstrøm, Paul Ragnar Berg and Monica Hongrø Solbakken for sampling in Sørøya.

## Availability of data and materials

The data set generated and analyzed for this study is available in the European Nucleotide Archive (ENA), study accession number PRJEB22384, http://www.ebi.ac.uk/ena/data/view/PRJEB29346.

## Supplementary file captions

**File S1: Multiple alignment of *Vibrionales* 16S V4 sequences in .fasta format.**

The file contains a .fasta-formatted multiple alignment of the 16S V4 region of the most highly abundant *Photobacterium, Vibrio* and *Aliivibrio* species (Table S4).

**File S2: *Lux* gene sequences of *Photobacterium kishitanii* in .fasta format.**

The file contains, for each gene in the lux operon, the reference genome sequence and the consensus sequence from both Lofoten and Sørøya.

## Supplementary table captions

**Table S1: Metadata**

Metadata collected for all specimens used in the study. Red and blue bars are applied to visualize associations between weight, length and age.

**Table S2: Filtering parameters**

Parameters used for quality filtering and trimming of metagenomic shotgun sequences in *Trimmomatic (*ver. 0.36) and *PRINSEQ-lite (*ver. 0.20.4).

**Table S3: *Vibrionales* genomes used for ANI-and variation analysis**

Overview of the 15 *Vibrionales* reference genomes used for genome similarity analysis measured by Average Nucleotide Identity (ANI) and variation analysis for the identification of single nucleotide variants (SNVs) within each genome. These genomes had the highest mean abundance among our samples after reference mapping.

**Table S4: Accessions used for 16S V4 sequence analysis**

The table shows all genomes (assemblies) used for the retrieval of 16S sequences (“..rna_from_genomic.fna.gz) used in the multiple alignment of *Vibrionales* V4 sequences.

**Table S5: Similarity (% identity) of V4 sequences**

The matrix shows the % identity between the 16S V4 region from the different *Vibrionales* species. Green cells indicate an identity of 100%, yellow an identity >= 97%.

**Table S6: *Vibrionales* reference genomes**

The table lists all reference genomes used for mapping of paired-end reads from the 19 intestinal microbiome samples.

**Table S7: Reference genomes similarity**

Results from genome similarity analyses based on average nucleotide identity (ANI) and mash distance. From the top: Table A) Jspecies website - ANIb, Table B) Jspecies website - Tetra, Table C) mash and Table D) fastANI. Plots in Fig. S3 are based on data from fastANI and mash.

**Table S8: Alpha diversity differences - Linear regression model**

Results from linear regression analysis used in testing for significant effects of location, sex or length on alpha diversity. The beyond optimal model including all covariates is presented here. The “top-down” strategy, selecting suitable covariates through t-tests, results in an “optimal” model with no covariates, indicating that neither location, sex or length have a significant effect on alpha diversity.

**Table S9: Homogeneity tests and PERMANOVA results for three datasets**

Results from homogeneity and PERMANOVA tests on the datasets based on order-and species-level read counts. All tests were performed on normalized data using the beta diversity measures Bray-Curtis dissimilarity and Jaccard index. *P*-values are marked in bold; values < 0.05 indicate statistical significance.

**Table S10: *Photobacterium kishitanii lux* operon**

The table shows accession number, length, position, and mean coverage per sample for each of the genes in the *Photobacterium kishitanii lux* operon. Gene sequence are given in the rightmost column.

**Table S11. Sample sizes**

Number of reads per sample before, during and after the trimming and filtering steps. The final two columns show the number of classified paired-end reads and its percentage of all paired-end reads. The lower table shows a summary of the data per location.

**Table S12. Classification of sequences by *Kaiju (*ver. 1.5.0)**

The table shows the numbers of total classified reads and number of reads classified at the species and order level per sample. A detailed overview of reads per order-level taxon per sample starts at column P. The lower table shows the first part of the same data, but at a relative scale. Sums and mean values are given in the green and yellow rows.

**Table S13: Alpha diversity values**

Alpha diversity estimates of the Atlantic cod intestinal microbial samples, calculated from non-normalized counts of reads classified at order level. See also fig. S1.

**Table S14: Significance tests of SNV distributions.**

For each of the 15 *Vibrionales* reference genomes, the tables show statistics for PCA of SNV distribution (Fig. S4, Fig. S5), including significance of two first PC axes (Tracy-Widom Statistic) and significance of between-group testing (Chi-square test). No Tracy-Widom *p*-values are significant after sequential Bonferroni correction (right table).

**Table S15: Coverage breadth of *P. iliopiscarium* and *P. kishitanii***

The tables show per sample the portion of each reference genome that is covered at a sequencing depth of at least 5X or 10X.

**Table S16: Genome-and gene level features of *P. iliopiscarium* and *P. kishitanii***

For each of the two most highly abundant *Vibrionales* species in the Atlantic cod gut, the table shows information on assembly, total number of genes, total number of annotated genes and the number of genes with zero coverage in the Atlantic cod samples.

**Table S17: Functional annotation of the 698 missing *Photobacterium kishitanii* genes** The table shows functional annotation data for each of the 698 genes absent in the *Photobacterium kishitanii* strain associated with Atlantic cod compared to its most closely related reference genome (GCF_000613045.2).

## Supplementary figure captions

**Figure S1:**
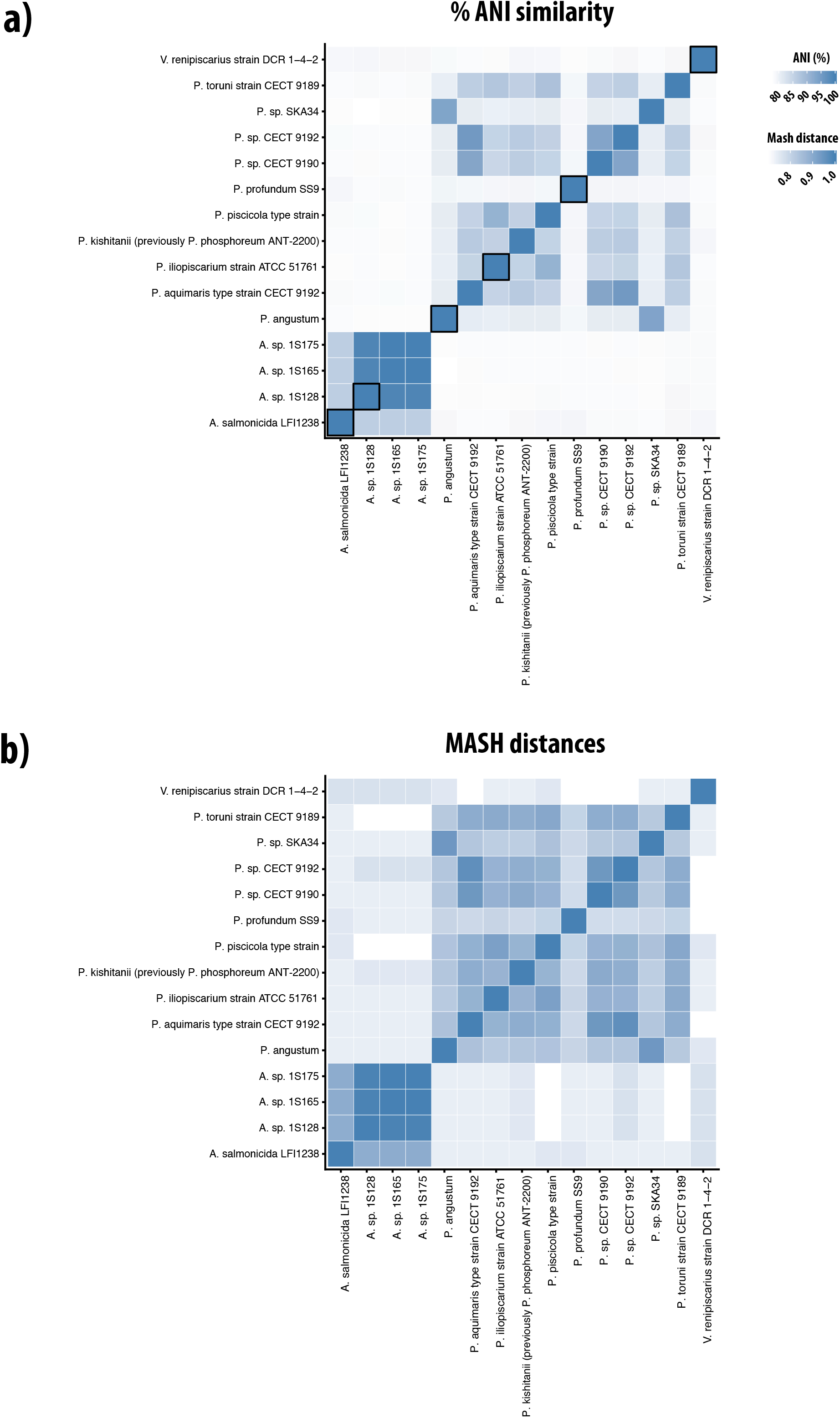
Similarity of 15 *Vibrionales* reference genomes. Heatmap showing the similarity of the reference genomes of the 15 most abundant *Vibrionales* species present in the Atlantic cod samples, based on (**A)** % Average Nucleotide Identity and (**B)** MASH distance. Squares with a black border in panel (A) represent the selection of refence genomes presented in Fig. 4.

**Figure S2:**
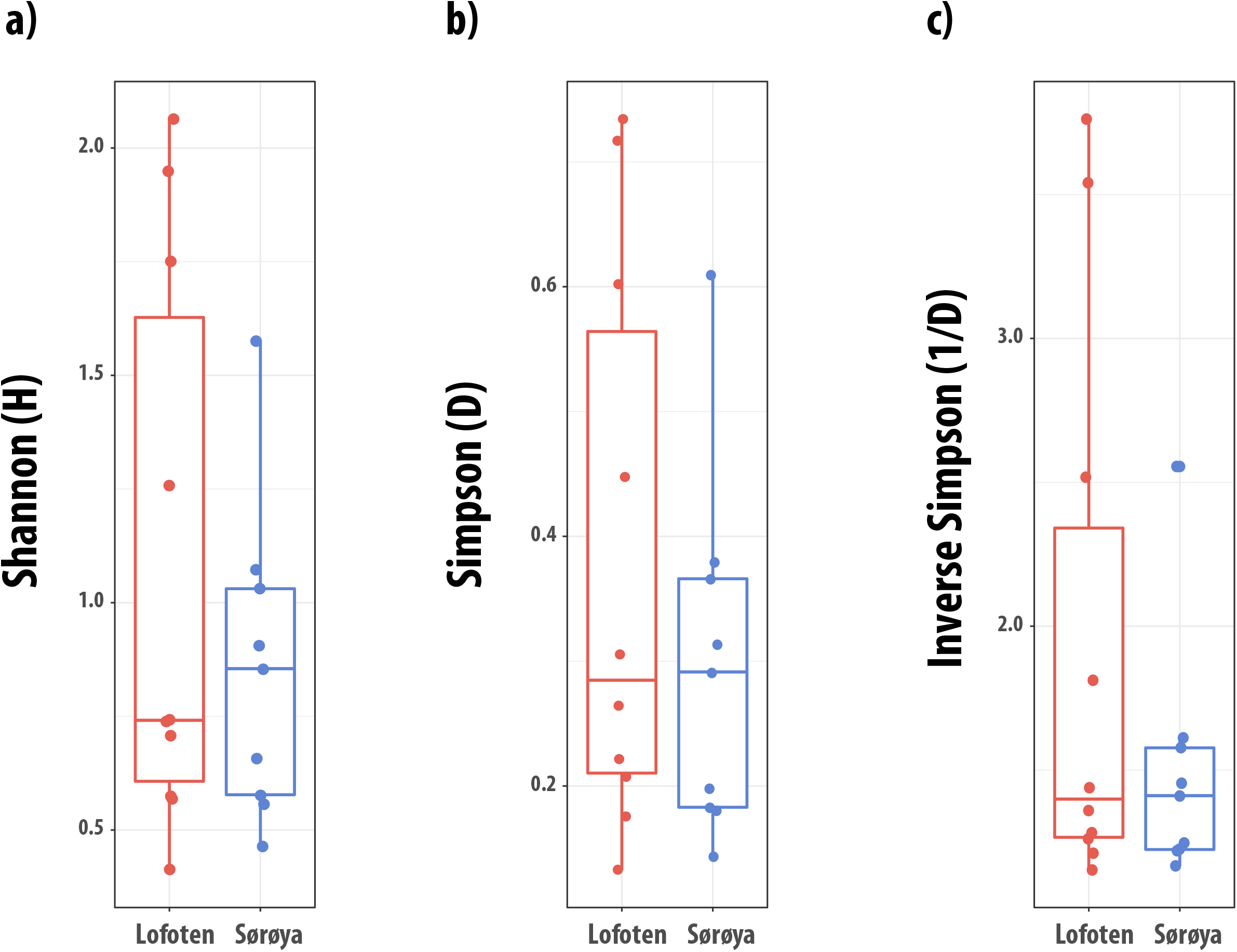
Within-sample diversity of Atlantic cod. Boxplots of Shannon (**A)**, Simpson (**B)** and Inverse Simpson (**C)** diversity in samples from Lofoten and Sørøya. The samples are grouped by location, and each of the 19 individuals is represented by a point. The middle band represents the median, while the upper and lower band shows the 75th and 25th percentile. The boxplots also show the minimum and maximum alpha diversity values.

**Figure S3:**
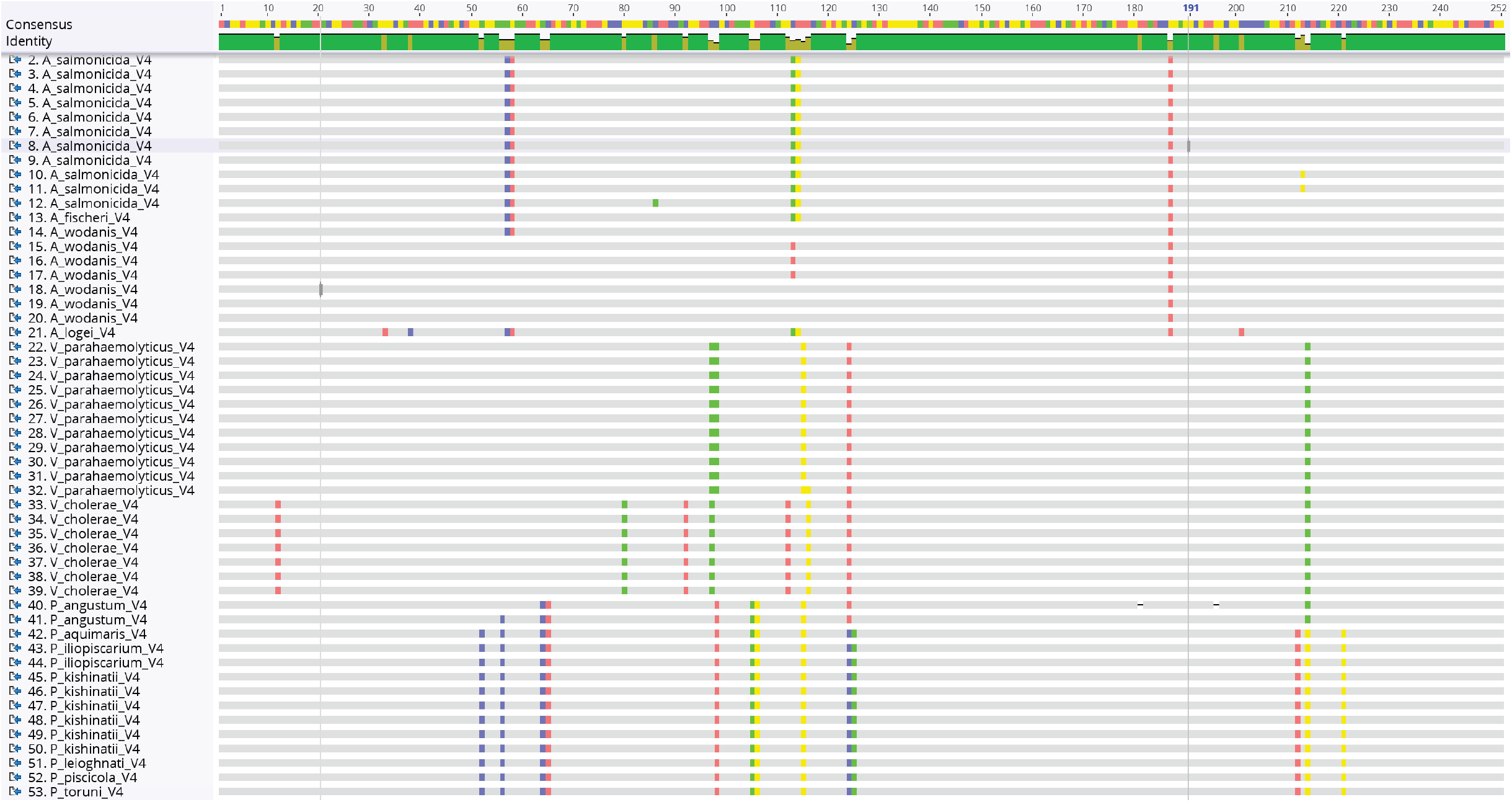
Multiple alignment of *Vibrionales* 16S V4 sequences. The figure shows a multiple alignment of the 16S V4 region of the most highly abundant *Photobacterium, Vibrio* and *Aliivibrio* species (Table S4). The full alignment is supplied as a .fasta file in File S1, and the corresponding similarity matrix is shown in Table S5.

**Figure S4:**
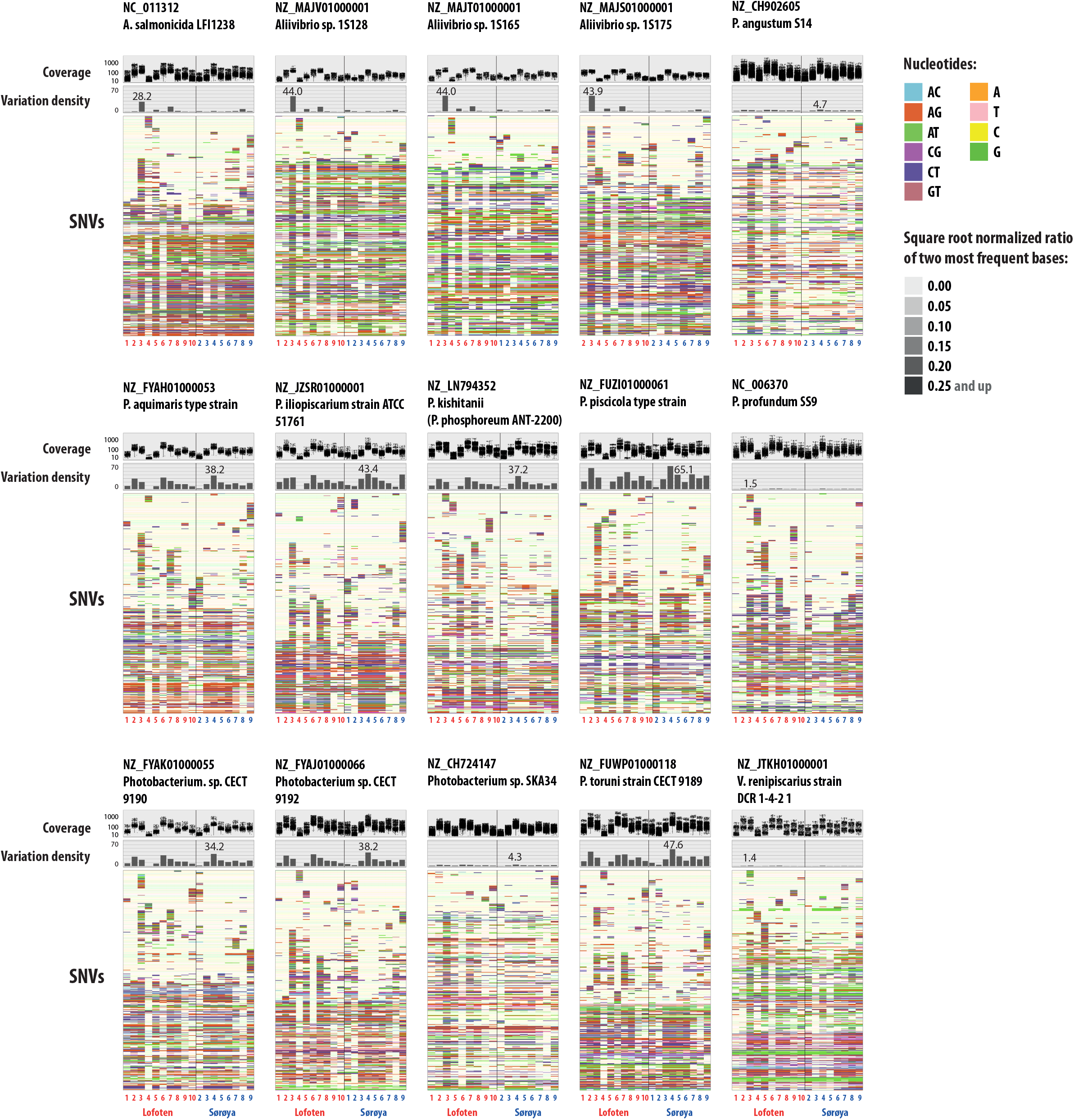
Variation analysis of 15 *Vibrionales* reference genomes. For each of the 15 most abundant *Vibrionales* genomes, the figure displays (from top to bottom), (1) read coverage per single nucleotide variant (SNV) position in each sample, (2) variation density (number of variable positions per Kbp. reported in each individual sample, independent of coverage in the other samples) per sample and (3) heatmap of a randomly chosen subset of 400 SNVs. In the heatmap, each row represents a unique variable nucleotide position, where the color of each tile represents the two most frequent competing nucleotides in that position. The shade of each tile represents the square root-normalized ratio of the most frequent two bases at that position (i.e., the more variation in a nucleotide position, the less pale the tile is). The y-axis of the coverage-and variation density plots are scaled across the reference genomes, and the mean SNV coverage across all samples is noted to the right of each coverage plot. For each genome, the density plot is annotated with the maximum variation density value. The total number of SNVs identified per reference genome is noted in parentheses after the strain name. Samples from Lofoten are numbered in red, while samples from Sørøya are numbered in blue.

**Figure S5:**
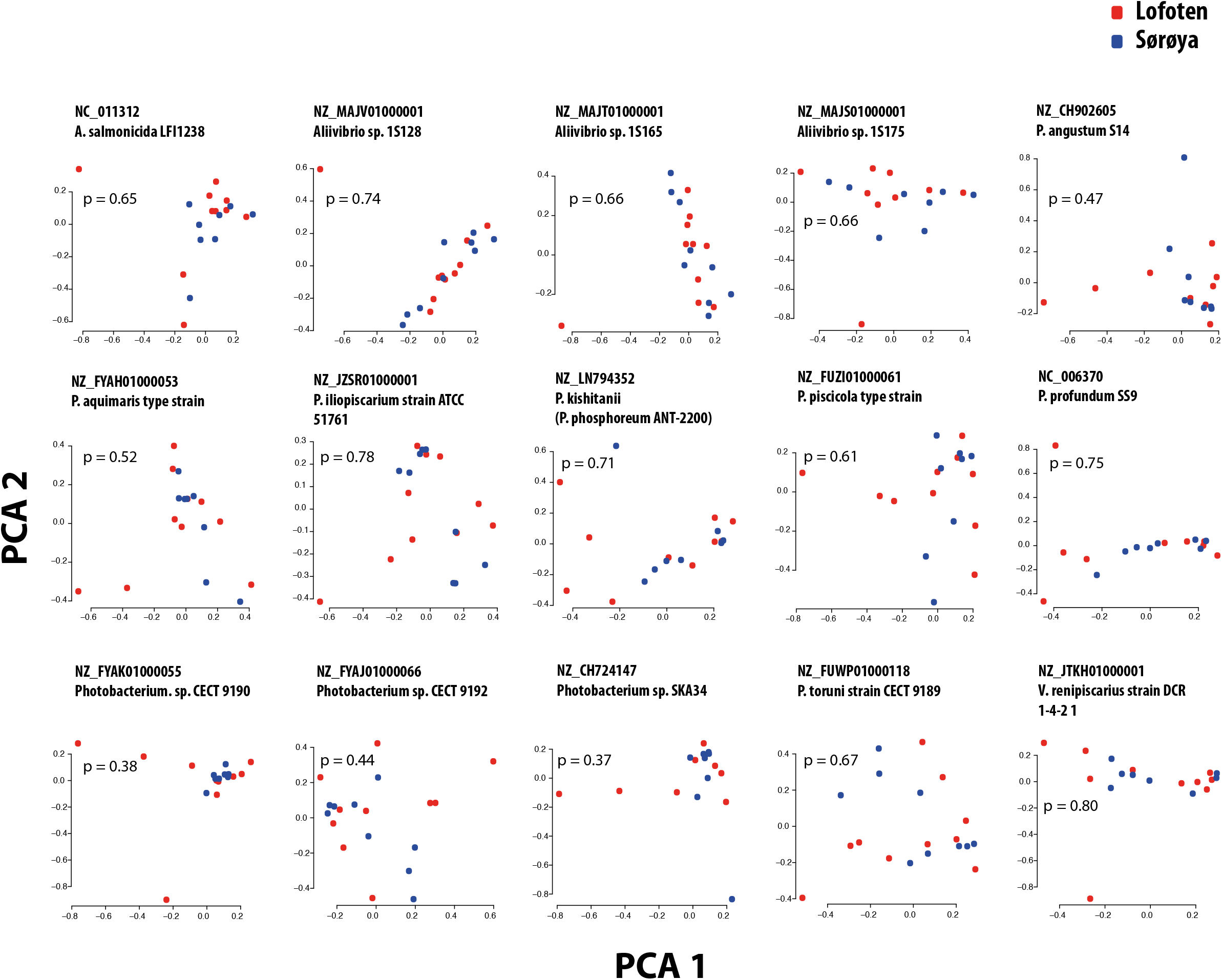
Principal component analysis (PCA) of SNVs in Lofoten and Sørøya. Principal component analysis of single nucleotide variant (SNV) distribution in individuals from Lofoten (red) and Sørøya (blue). Each plot represents one of the 15 *Vibrionales* reference genomes with highest mean abundance, and the ordering is similar to the 15 SNV plots in Fig. S4. Overlapping clusters indicate no spatial separation of the Atlantic cod intestinal microbiome. The *p*-value from a Chi-squared test of geographical differences is included in each plot. Detailed statistics for each plot are given in Table S14.

**Figure S6:**
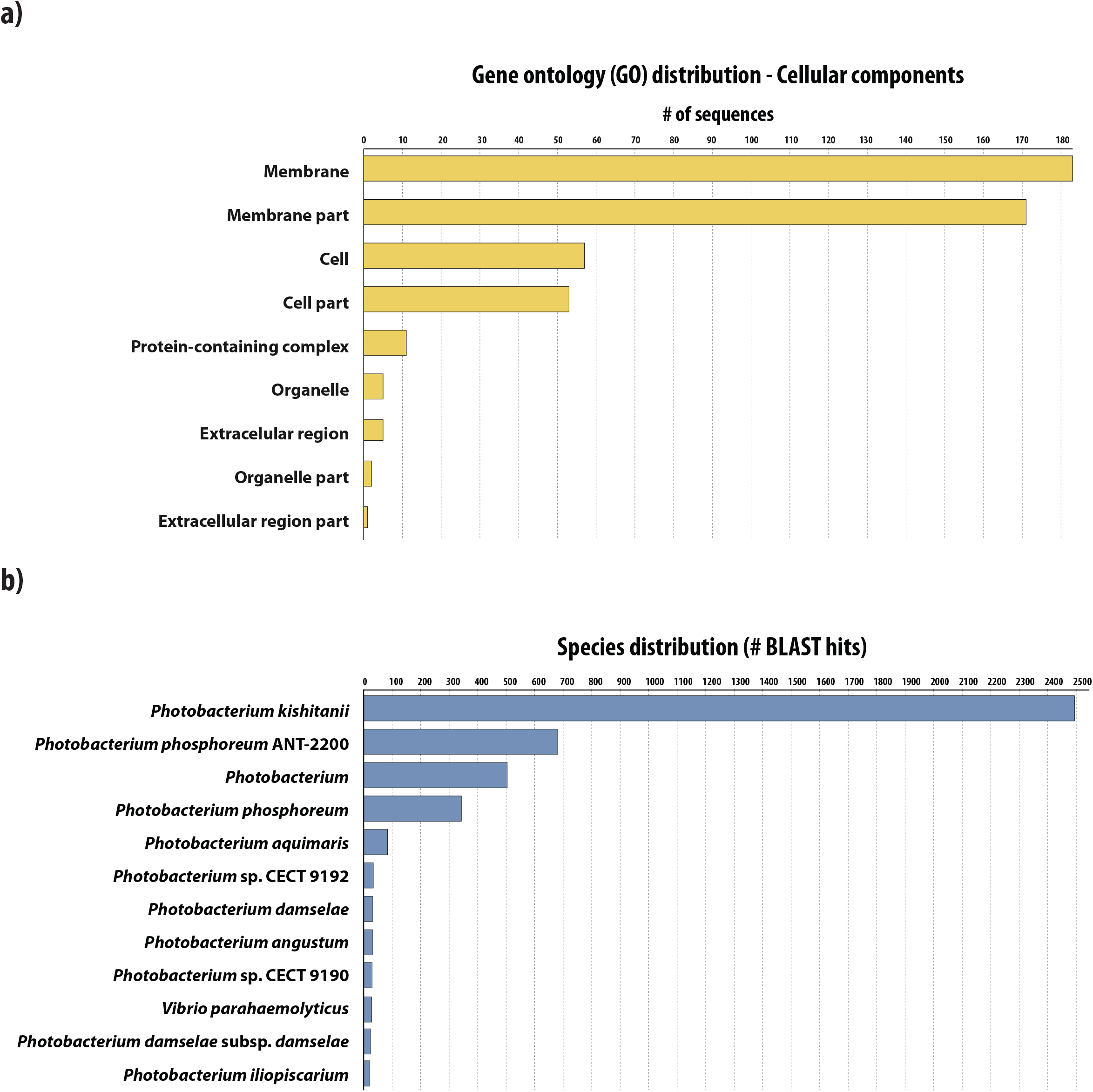
Functional analysis of the 698 *P. kishitanii* genes missing in our closely related *Photobacterium* strain. **(A)** Overview of the complexes or compartments where the gene products of the missing genes are potentially active. The x-axis represents the number of gene sequences assigned to each GO (Gene ontology) category. (**B)** Overview of the taxonomy affiliated with hits from a blast search with the 698 gene sequences. The top five hits were kept for each individual sequence search. The x-axis represents the number of hits associated with each taxonomical category.

**Figure S7:**
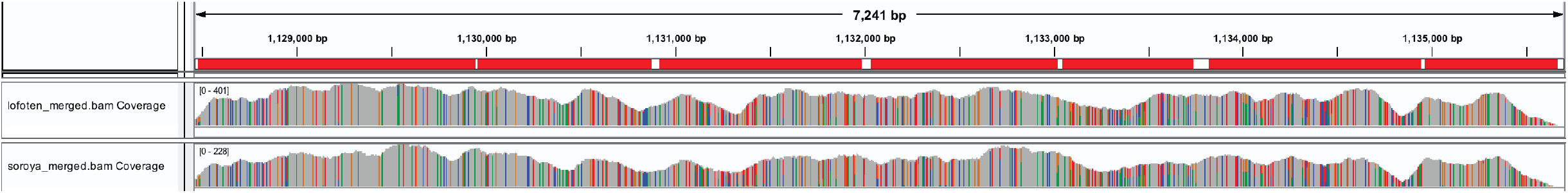
Coverage of the *lux* operon in *Photobacterium kishitanii*. The figure shows the coverage of the *Photobacterium kishitanii lux* operon, as displayed in the *Integrative Genomics Viewer*. The coverage of individual samples has been summed for both Lofoten (max. coverage: 401X) and Sørøya (max. coverage 228X). The horizontal red bars represent the individual *lux* genes in the operon (*luxC, luxD, luxA, luxB, luxF, luxE* and *luxG*).

## References

Altschul, S.F., Gish, W., Miller, W., Myers, E.W., and Lipman, D.J. (1990) Basic Local Alignment Search Tool. J. Mol. Biol. 215: 403–410.

Amore, R.D., Ijaz, U.Z., Schirmer, M., Kenny, J.G., Gregory, R., Darby, A.C., et al. (2016) A comprehensive benchmarking study of protocols and sequencing platforms for 16S rRNA community profiling. BMC Genomics 17:.

Andrews, C.C., Karl, D.M., Small, L.F., and Fowler, S.W. (1984) Metabolic activity and bioluminescence of oceanic faecal pellets and sediment trap particles. Nature 307: 539.

Andrews, S. (2010) FastQC. Babraham Bioinformatics.

Arndt, D., Grant, J.R., Marcu, A., Sajed, T., Pon, A., Liang, Y., and Wishart, D.S. (2016) PHASTER: a better, faster version of the PHAST phage search tool. Nucleic Acids Res. 44: 16–21.

Artimo, P., Jonnalagedda, M., Arnold, K., Baratin, D., Flegel, V., Fortier, A., et al. (2012) ExPASy: SIB bioinformatics resource portal. Nucleic Acids Res. 40: 597–603.

Ast, J.C. and Dunlap, P. V. (2005) Phylogenetic resolution and habitat specificity of members of the Photobacterium phosphoreum species group. Environ. Microbiol. 7: 1641–1654.

Bairagi, A., Ghosh, K.S., Kumar, S., and Ray, A.K. (2002) Enzyme producing bacterial flora isolated from fish. Aquac. Int. 10: 109–121.

Benjamini, Y. and Speed, T.P. (2012) Summarizing and correcting the GC content bias in high-throughput sequencing. Nucleic Acids Res. 40: 1–14.

Birtel, J., Walser, J., Pichon, S., and Bürgmann, H. (2015) Estimating Bacterial Diversity for Ecological Studies: Methods, Metrics, and Assumptions. PLoS One 1–23.

Bolger, A.M., Lohse, M., and Usadel, B. (2014) Trimmomatic: A flexible trimmer for Illumina sequence data. Bioinformatics 30: 2114–2120.

van den Boogaart, K.G. and Tolosana-Delgado, R. (2013) Analyzing Compositional Data with R. Springer Verlag, Heidelberg.

van den Boogaart, K.G. and Tolosana-Delgado, R. (2008) “compositions”: A unified R package to analyze compositional data. Comput. Geosci. 37: 320–338.

Caipang, C.M.A., Brinchmann, M.F., and Kiron, V. (2010) Antagonistic activity of bacterial isolates from intestinal microbiota of Atlantic cod, Gadus morhua, and an investigation of their immunomodulatory capabilities. Aquac. Res. 41: 249–256.

Camacho, C., Coulouris, G., Avagyan, V., Ma, N., Papadopoulos, J., Bealer, K., and Madden, T.L. (2009) BLAST+: architecture and applications. BMC Bioinformatics 9: 1–9.

Cohen, D.M., Inada, T., Iwamoto, T., and Scialabba, N. (1990) Gadiform fishes of the world (Order Gadiformes). An annotated and illustrated catalogue of cods, hakes, grenadiers and other gadiform fishes known to date. In, FAO Species Catalogue.

Colston, T.J. and Jackson, C.R. (2016) Microbiome evolution along divergent branches of the vertebrate tree of life: what is known and unknown. Mol. Ecol. 25: 3776–3800.

Conesa, A. and Götz, S. (2008) Blast2GO: A Comprehensive Suite for Functional Analysis in. Int. J. Plant Genomics.

Conesa, A., Götz, S., García-gómez, J.M., Terol, J., Talón, M., Genómica, D., et al. (2005) Blast2GO: a universal tool for annotation, visualization and analysis in functional genomics research. Bioinformatics 21: 3674–3676.

Dehler, C.E., Secombes, C.J., and Martin, S.A.M. (2017) Environmental and physiological factors shape the gut microbiota of Atlantic salmon parr (Salmo salar L.). Aquaculture 467: 149–157.

Desai, A.R., Links, M.G., Collins, S.A., Mansfield, G.S., Drew, M.D., Van Kessel, A.G., and Hill, J.E. (2012) Effects of plant-based diets on the distal gut microbiome of rainbow trout (Oncorhynchus mykiss). Aquaculture 350–353: 134–142.

Dhanasiri, A.K.S., Brunvold, L., Brinchmann, M.F., Korsnes, K., Bergh, Ø., and Kiron, V. (2011) Changes in the intestinal microbiota of wild Atlantic cod (Gadus morhua L.) upon captive rearing. Microb. Ecol. 61: 20–30.

ecogenomic.org (2013) Cetobacterium somerae - GC content. gtdb.ecogenomic.org.

Egerton, S., Culloty, S., Whooley, J., Stanton, C., and Ross, R.P. (2018) The gut microbiota of marine fish. Front. Microbiol. 9: 1–17.

Eren, A.M., Esen, C., Quince, C., Vineis, J.H., Morrison, H.G., Sogin, M.L., and Delmont, T.O. (2015a) Anvi’o: An advanced analysis and visualization platform for ‘omics data. PeerJ 3: 1–29.

Eren, A.M., Esen, C., Quince, C., Vineis, J.H., Morrison, H.G., Sogin, M.L., and Delmont, T.O. (2015b) Visualizing SNV profiles using R.

Froese, Rainer and Pauly, D. (2012) Species Fact Sheets: Gadus morhua (Linnaeus, 1758). Geneious ver. 10.2.2, Geneious.

Genome Reference Consortium (2009) Genome Reference Consortium Human Build 37 (GRCh37).

Ghanbari, M., Kneifel, W., and Domig, K.J. (2015) A new view of the fish gut microbiome: Advances from next-generation sequencing. Aquaculture 448: 464–475.

Gilbert, J.A., Meyer, F., Jansson, J., Gordon, J., Pace, N., Ley, R., et al. (2010) The Earth Microbiome Project: Meeting report of the “1st EMP meeting on sample selection and acquisition” at Argonne National Laboratory October 6th 2010.

Givens, C.E., Ransom, B., Bano, N., and Hollibaugh, J.T. (2015) Comparison of the gut microbiomes of 12 bony fish and 3 shark species. Mar. Ecol. Prog. Ser. 518: 209–223.

Godø, O.R. and Michalsen, K. (2000) Migratory behaviour of North-east Arctic cod, studied by use of data storage tags. Fish. Res. 48: 127–140.

Götz, S., Arnold, R., Sebastián-león, P., Martín-rodríguez, S., Tischler, P., Jehl, M., et al. (2011) B2G-FAR, a species-centered GO annotation repository. Bioinformatics 27: 919–924.

Götz, S., Garcia-Gomez, J.M., Terol, J., Williams, T.D., Nagaraj, S.H., Nueda, M.J., et al. (2008) High-throughput functional annotation and data mining with the Blast2GO suite. Nucleic Acids Res. 36: 3420–3435.

Hamre, K. (2006) Nutrition in cod (Gadus morhua) larvae and juveniles. ICES J. Mar. Sci. 63: 267–274.

Hennersdorf, P., Mrotzek, G., Abdul-Aziz, M.A., and Saluz, H.P. (2016) Metagenomic analysis between free-living and cultured Epinephelus fuscoguttatus under different environmental conditions in Indonesian waters. Mar. Pollut. Bull. 110: 726–734.

Holm, S. (1979) A Simple Sequentially Rejective Multiple Test Procedure. Scand. J. Stat. 6: 65–70.

Izvekova, G., Izvekov, E., and Plotnikov, A. (2007) Symbiotic microflora in fishes of different ecological groups. Biol. Bull. 34: 610–618.

Jain, C., Rodriguez-R, L.M., Phillippy, A.M., Konstantinidis, K.T., and Aluru, S. (2018) High throughput ANI analysis of 90K prokaryotic genomes reveals clear species boundaries. Nat. Commun. 9: 1–8. Joint Genome Institute BBMap.

Jones, P., Binns, D., Chang, H., Fraser, M., Li, W., Mcanulla, C., et al. (2014) Sequence analysis InterProScan 5: genome–scale protein function classification. Bioinformatics 30: 1236–1240.

Kim, D.-H., Brunt, J., and Austin, B. (2007) Microbial diversity of intestinal contents and mucus in rainbow trout (Oncorhynchus mykiss). J. Appl. Microbiol. 102: 1654–64.

Konstantinidis, K.T., Ramette, A., and Tiedje, J.M. (2006) The bacterial species definition in the genomic era. Philos. Trans. R. Soc. B Biol. Sci. 361: 1929–1940.

Köster, J. and Rahmann, S. (2012) Snakemake - a scalable bioinformatics workflow engine. Bioinformatics 28: 2520–2522.

Li, H. (2012) Seqtk.

Li, H. and Durbin, R. (2009) Fast and accurate short read alignment with Burrows – Wheeler transform. Bioinformatics 25: 1754–1760.

Li, H., Handsaker, B., Wysoker, A., Fennell, T., Ruan, J., Homer, N., et al. (2009) The Sequence Alignment/Map format and SAMtools. Bioinformatics 25: 2078–2079.

Lie, K.K., Tørresen, O.K., Solbakken, M.H., Rønnestad, I., Tooming-klunderud, A., Nederbragt, A.J., et al. (2018) Loss of stomach, loss of appetite? Sequencing of the ballan wrasse (Labrus bergylta) genome and intestinal transcriptomic profiling illuminate the evolution of loss of stomach function in fish. BMC Genomics 19: 1–17.

Link, J.S., Bogstad, B., Sparholt, H., and Lilly, G.R. (2009) Trophic role of Atlantic cod in the ecosystem. Fish Fish. 10: 58–87.

Liu, Z., Desantis, T.Z., Andersen, G.L., and Knight, R. (2008) Accurate taxonomy assignments from 16S rRNA sequences produced by highly parallel pyrosequencers. Nucleic Acids Res. 36: 1–11.

Llewellyn, M.S., Boutin, S., Hoseinifar, S.H., and Derome, N. (2014) Teleost microbiomes: the state of the art in their characterization, manipulation and importance in aquaculture and fisheries. Front. Microbiol. 5: 207.

Llewellyn, M.S., McGinnity, P., Dionne, M., Letourneau, J., Thonier, F., Carvalho, G.R., et al. (2016) The biogeography of the Atlantic salmon (Salmo salar) gut microbiome. ISME J 10: 1280–1284.

MacDonald, N.L., Stark, J.R., and Austin, B. (1986) Bacterial microflora in the gastro-intestinal tract of Dover sole (Solea solea L.), with emphasis on the possible role of bacteria in the nutrition of the host. FEMS Microbiol. Lett. 35: 107–111.

Machado, H. and Gram, L. (2017) Comparative genomics reveals high genomic diversity in the genus Photobacterium. Front. Microbiol. 8: 1–14.

Machado, H. and Gram, L. (2015) The fur gene as a new phylogenetic marker for Vibrionaceae species identification. Appl. Environ. Microbiol. 81: 2745–2752.

Malmstrøm, M., Matschiner, M., Tørresen, O.K., Star, B., Snipen, L.G., Hansen, T.F., et al. (2016) Evolution of the immune system influences speciation rates in teleost fishes. Nat. Genet. 48: 1204–1210.

Martin-Antonio, B., Manchado, M., Infante, C., Zerolo, R., Labella, A., Alonso, C., and Borrego, J.J. (2007) Intestinal microbiota variation in Senegalese sole (Solea senegalensis) under different feeding regimes. Aquac. Res. 38: 1213–1222.

McMurdie, P.J. and Holmes, S. (2014) Waste Not, Want Not: Why Rarefying Microbiome Data Is Inadmissible. PLoS Comput. Biol. 10:.

Menzel, P. and Krogh, A. (2016) Fast and sensitive taxonomic classification for metagenomics with Kaiju. Nat. Commun. 7:.

Merrifield, D.L. and Rodiles, A. (2015) The fish microbiome and its interactions with mucosal tissues. In, Mucosal health in Aquaculture.

Michalsen, K., Johannesen, E., and Bogstad, B. (2008) Feeding of mature cod (Gadus morhua) on the spawning grounds in Lofoten. ICES J. Mar. Sci. 65: 571–580.

Mitchell, A.L., Attwood, T.K., Babbitt, P.C., Blum, M., Bork, P., Bridge, A., et al. (2019) InterPro in 2019: improving coverage, classification and access to protein sequence annotations. 1–10.

Noecker, C., Mcnally, C.P., Eng, A., and Borenstein, E. (2016) High-resolution characterization of the human microbiome. Transl. Res. 179: 7–23.

Norecopa Norecopa guidelines for animal experiments.

O’Leary, N.A., Wright, M.W., Brister, J.R., Ciufo, S., Haddad, D., McVeigh, R., et al. (2016) Reference sequence (RefSeq) database at NCBI: Current status, taxonomic expansion, and functional annotation. Nucleic Acids Res. 44: D733–D745.

Oksanen, J., Blanchet, F.G., Friendly, M., Kindt, R., Legendre, P., McGlinn, D., et al. (2017) vegan: Community Ecology Package R package version 2.4–3.

Onarheim, A.M. and Raa, J. (1990) Characteristics and possible biological significance of an autochthonous flora in the intestinal mucosa of sea-water fish. Syst. Appl. Microbiol. 17: 197–201.

Onarheim, A.M., Wiik, R., Burghardt, J., and Stackebrandt, E. (1994) Characterization & identification of two Vibrio species indigenous to the intestine of fish in cold sea water; description of Vibrio iliopiscarious sp. nov. Syst. Appl. Microbiol. 17: 370–379.

Ondov, B.D., Treangen, T.J., Melsted, P., Mallonee, A.B., Bergman, N.H., Koren, S., and Phillippy, A.M. (2016) Mash: Fast genome and metagenome distance estimation using MinHash. Genome Biol. 17: 1–14.

Patterson, N., Price, A.L., and Reich, D. (2006) Population Structure and Eigenanalysis. PLoS Genet. 2:.

R Core Team (2017) R: A language and environment for statistical computing. R Found. Stat. Comput. Vienna, Austria.

Racine, J.S. (2010) Rstudio: A platform-independent ide for R and SWEAVE. Financ. Dev. 47: 36–37.

Ranjan, R., Rani, A., Metwally, A., McGee, H.S., and Perkins, D.L. (2016) Analysis of the microbiome: Advantages of whole genome shotgun versus 16S amplicon sequencing. Biochem. Biophys. Res. Commun. 469: 967–977.

Rawls, J.F., Samuel, B.S., and Gordon, J.I. (2004) Gnotobiotic zebrafish reveal evolutionarily conserved responses to the gut microbiota. Proc. Natl. Acad. Sci. 101: 4596–4601.

Ray, A.K., Ghosh, K., and Ringø, E. (2012) Enzyme-producing bacteria isolated from fish gut: A review. Aquac. Nutr. 18: 465–492.

Righton, D.A., Andersen, K.H., Neat, F., Thorsteinsson, V., Steingrund, P., Svedäng, H., et al. (2010) Thermal niche of Atlantic cod (Gadus morhua): Limits, tolerance and optima. Mar. Ecol. Prog. Ser. 420: 1–13.

Riiser, E.S., Haverkamp, T.H.A., Borgan, Ø., Jakobsen, K.S., Jentoft, S., and Star, B. (2018) A single Vibrionales 16S rRNA oligotype dominates the intestinal microbiome in two geographically separated Atlantic cod populations. Front. Microbiol. 9: 1–14.

Ringø, E., Sperstad, S., Myklebust, R., Refstie, S., and Krogdahl, Å. (2006) Characterisation of the microbiota associated with intestine of Atlantic cod (Gadus morhua L.). Aquaculture 261: 829–841.

Robinson, J.T., Thorvaldsdóttir, H., Winckler, W., Guttman, M., Lander, E.S., Getz, G., and Mesirov., J.P. (2011) Integrative genomics viewer. Nat. Biotechnol. 29: 24–26.

Roeselers, G., Mittge, E.K., Stephens, W.Z., Parichy, D.M., Cavanaugh, C.M., Guillemin, K., and Rawls, J.F. (2011) Evidence for a core gut microbiota in the zebrafish. ISME J. 5: 1595–608.

Romero, J., Ringø, E., and Merrifield, D.L. (2014) The Gut Microbiota of Fish. Aquac. Nutr. 75–100.

Ruby, E.G. and Morin, J. (1979) Luminous Enteric Bacteria of Marine Fishes: a Study of Their Distribution, Densities, and Dispersion. Appl. Environ. Microbiol. 38: 406–411.

Samuelsen, O.B., Nerland, A.H., Jørgensen, T., Schrøder, M.B., Svåsand, T., and Bergh, Ø. (2006) Viral and bacterial diseases of Atlantic cod Gadus morhua, their prophylaxis and treatment: a review. Dis. Aquat. Organ. 71: 239–254.

Sawabe, T., Kita-Tsukamoto, K., and Thompson, F.L. (2007) Inferring the evolutionary history of Vibrios by means of multilocus sequence analysis. J. Bacteriol. 189: 7932–7936.

Schmidt, V., Amaral-Zettler, L., Davidson, J., Summerfelt, S., and Good, C. (2016) Influence of fishmeal-free diets on microbial communities in Atlantic salmon (Salmo salar) recirculation aquaculture systems. Appl. Environ. Microbiol. 82: 4470–4481.

Schmieder, R. and Edwards, R. (2011) Quality control and preprocessing of metagenomic datasets. Bioinformatics 27: 863–864.

Shakya, M., Quince, C., Campbell, J.H., Yang, Z.K., Schadt, C.W., and Podar, M. (2013) Comparative metagenomic and rRNA microbial diversity characterization using archaeal and bacterial synthetic communities. Environ. Microbiol. 15: 1882–1899.

Smith, C.J., Danilowicz, B.S., and Meijer, W.G. (2007) Characterization of the bacterial community associated with the surface and mucus layer of whiting (Merlangius merlangus). FEMS Microbiol. Ecol. 62: 90–97.

Solbakken, M.H., Tørresen, O.K., Nederbragt, A.J., Seppola, M., Gregers, T.F., Jakobsen, K.S., and Jentoft, S. (2016) Evolutionary redesign of the Atlantic cod (Gadus morhua L.) Toll-like receptor repertoire by gene losses and expansions. Sci. Rep. 6: 1–14.

Star, B., Haverkamp, T.H., Jentoft, S., and Jakobsen, K.S. (2013) Next generation sequencing shows high variation of the intestinal microbial species composition in Atlantic cod caught at a single location. BMC Microbiol. 13: 248.

Star, B. and Jentoft, S. (2012) Why does the immune system of Atlantic cod lack MHC II? Bioessays 34: 648–51.

Star, B., Nederbragt, A.J., Jentoft, S., Grimholt, U., Malmstrøm, M., Gregers, T.F., et al. (2011) The genome sequence of Atlantic cod reveals a unique immune system. Nature 477: 207–10.

Sugita, H., Kawasaki, J., and Deguchi, Y. (1997) Production of amylase by the intestinal microflora in cultured freshwater fish. Lett. AppliedMicrobiology 1 24: 105–108.

Sugita, H., Miyajima, C., and Deguchi, Y. (1991) The vitamin B12-producing ability of the intestinal microflora of freshwater fish. Aquaculture 92: 267–276.

Sullam, K.E., Essinger, S.D., Lozupone, C.A., O’Connor, M.P., Rosen, G.L., Knight, R., et al. (2012) Environmental and ecological factors that shape the gut bacterial communities of fish: a meta-analysis. Mol. Ecol. 21: 3363–78.

Takemura, A.F., Chien, D.M., and Polz, M.F. (2014) Associations and dynamics of Vibrionaceae in the environment, from the genus to the population level. Front. Microbiol. 5:.

Talwar, C., Nagar, S., Lal, R., and Negi, R.K. (2018) Fish Gut Microbiome: Current Approaches and Future Perspectives. Indian J. Microbiol.

Tarnecki, A.M., Burgos, F.A., Ray, C.L., and Arias, C.R. (2017) Fish intestinal microbiome: diversity and symbiosis unravelled by metagenomics. J. Appl. Microbiol.

Thorvaldsdóttir, H., Robinson, J.T., and Mesirov, J.P. (2013) Integrative Genomics Viewer (IGV): high-performance genomics data visualization and exploration. Brief. Bioinforma. 14: 178–192.

Tørresen, O.K., Star, B., Jentoft, S., Reinar, W.B., Grove, H., Miller, J.R., et al. (2017) An improved genome assembly uncovers prolific tandem repeats in Atlantic cod. BMC Genomics 18: 1–23.

Tsuchiya, C., Sakata, T., and Sugita, H. (2008) Novel ecological niche of Cetobacterium somerae, an anaerobic bacterium in the intestinal tracts of freshwater fish. Lett. Appl. Microbiol. 46: 43–48.

Tyagi, A., Singh, B., Billekallu, N.K., and Niraj, T. (2019) Shotgun metagenomics offers novel insights into taxonomic compositions, metabolic pathways and antibiotic resistance genes in fish gut microbiome. Arch. Microbiol.

Uchii, K., Matsui, K., Yonekura, R., Tani, K., Kenzaka, T., Nasu, M., and Kawabata, Z. (2006) Genetic and physiological characterization of the intestinal bacterial microbiota of Bluegill (Lepomis macrochirus) with three different feeding habits. Microb. Ecol. 51: 277–284.

Urakawa, H., Kita-Tsukamoto, K., and Ohwada, K. (1999) Reassessment of the taxonomic position of Vibrio iliopiscarium (Onarheim et al. 1994) and proposal for Photobacterium iliopiscarium comb. nov. Int. J. Syst. Bacteriol. 49: 257–260.

Urbanczyk, H., Ast, J.C., and Dunlap, P. V. (2011) Phylogeny, genomics, and symbiosis of Photobacterium. FEMS Microbiol. Rev. 35: 324–342.

Valdenegro-Vega, V., Naeem, S., Carson, J., Bowman, J.P., Tejedor del Real, J.L., and Nowak, B. (2013) Culturable microbiota of ranched southern bluefin tuna (Thunnus maccoyii Castelnau). J. Appl. Microbiol. 115: 923–932.

Vasileiadis, S., Puglisi, E., Arena, M., Cappa, F., Cocconcelli, P.S., and Trevisan, M. (2012) Soil Bacterial Diversity Screening Using Single 16S rRNA Gene V Regions Coupled with Multi-Million Read Generating Sequencing Technologies. PLoS One 7:.

Verner-Jeffreys, D.W., Shields, R., Bricknell, I.R., and Birkbeck, T.H. (2003) Changes in the gut-associated microflora during the development of Atlantic halibut (Hippoglossus rhippoglossus L.) larvae in three British hatcheries. Aquaculture 219: 21–42.

Wang, A.R., Ran, C., Ringø, E., and Zhou, Z.G. (2017) Progress in fish gastrointestinal microbiota research. Rev. Aquac. 1–15.

Ward, N.L., Steven, B., Penn, K., Methé, B.A., and Detrich, W.H. (2009) Characterization of the intestinal microbiota of two Antarctic notothenioid fish species. Extremophiles 13: 679–685.

Wickham, H. (2009) Ggplot2.

Wu, S.G., Tian, J.Y., Gatesoupe, F.J., Li, W.X., Zou, H., Yang, B.J., and Wang, G.T. (2013) Intestinal microbiota of Gibel carp (Carassius auratus gibelio) and its origin as revealed by 454 pyrosequencing. World J. Microbiol. Biotechnol. 29: 1585–1595.

Xia, J.H., Lin, G., Fu, G.H., Wan, Z.Y., Lee, M., Wang, L., et al. (2014) The intestinal microbiome of fish under starvation. BMC Genomics 15: 266.

Xing, M., Hou, Z., Yuan, J., Liu, Y., Qu, Y., and Liu, B. (2013) Taxonomic and functional metagenomic profiling of gastrointestinal tract microbiome of the farmed adult turbot (Scophthalmus maximus). FEMS Microbiol. Ecol. 86: 432–443.

Ye, L., Amberg, J., Chapman, D., Gaikowski, M., and Liu, W.T. (2014) Fish gut microbiota analysis differentiates physiology and behavior of invasive Asian carp and indigenous American fish. ISME J. 8: 541–551.

Youssef, N., Sheik, C.S., Krumholz, L.R., Najar, F.Z., Roe, B.A., and Elshahed, M.S. (2009) Comparison of Species Richness Estimates Obtained Using Nearly Complete Fragments and Simulated Pyrosequencing-Generated Fragments in 16S rRNA Gene-Based Environmental Surveys. Appl. Environ. Microbiol. 75: 5227–5236.

Zarkasi, K.Z., Abell, G.C.J., Taylor, R.S., Neuman, C., Hatje, E., Tamplin, M.L., et al. (2014) Pyrosequencing-based characterization of gastrointestinal bacteria of Atlantic salmon (Salmo salar L.) within a commercial mariculture system. J. Appl. Microbiol. 117: 18–27.

Zarkasi, K.Z., Taylor, R.S., Abell, G.C.J., Tamplin, M.L., Glencross, B.D., and Bowman, J.P. (2016) Atlantic Salmon (Salmo salar L.) Gastrointestinal Microbial Community Dynamics in Relation to Digesta Properties and Diet. Microb. Ecol. 71: 589–603.

Zarubin, M., Belkin, S., Ionescu, M., and Genin, A. (2012) Bacterial bioluminescence as a lure for marine zooplankton and fish. Proc. Natl. Acad. Sci. 109: 853–857.

Zhang, J., Ding, X., Guan, R., Zhu, C., Xu, C., Zhu, B., et al. (2018) Evaluation of different 16S rRNA gene V regions for exploring bacterial diversity in a eutrophic freshwater lake. Sci. Total Environ. 618: 1254–1267.

Zhou, Y., Liang, Y., Lynch, K.H., Dennis, J.J., and Wishart, D.S. (2011) PHAST: A Fast Phage Search Tool. Nucleic Acids Res. 39: 347–352.

